# Species-specific epigenetic responses to drought stress of two sympatric oak species reflect their ecological preferences

**DOI:** 10.1101/2023.09.26.559529

**Authors:** B Rubio, G Le Provost, B Brachi, T Gerardin, O Brendel, J. Tost, Christian Daviaud, P Gallusci

## Abstract

- In a context of climate change, it is necessary to decipher the strategies established by plants to cope with limited water supply.
- Transcriptome, methylome and small RNA data were generated for two oak species with contrasting levels of drought tolerance (*Quercus robur* and *Quercus petraea*), under control and drought stress conditions
- All data are in line with a species-specific response to drought stress consistent with their ecological preferences. The biological processes associated with genomic regions identified in all datasets were mainly associated with parietal processes in *Q. petraea,* which may explain in part its better tolerance to water deprivation.
- A significant proportion of DNA methylation differences observed in control conditions between the two oak species were maintained during DS which may constitute a pool of epigenetic markers discriminating these two oak species. These markers were enriched in highly differentiated SNPs suggesting that some of them may be associated both with the ecological differences or intrinsic barriers to reproduction between the two species.
- An integrative approach of the three datasets revealed genomic co-locations of potential importance for forest three adaptation to drought stress.

## Introduction

European white oaks which belong to the Fagaceae family, are represented principally by two widely distributed species: the pedunculate (*Quercus robur* L., hereafter PO) and sessile (*Quercus petraea* (Matt.) Liebl., hereafter SO) oaks (Epron & Dreyer, 1990; Haneca *et al*., 2009). The distributions of PO and SO are determined principally by environmental factors (Epron & Dreyer, 1990; Haneca *et al*., 2009). PO, which can tolerate high levels of waterlogging, is found at mesic sites, whereas SO, a drought-tolerant species grows essentially in well-drained soils (Schmull & Thomas, 2000). PO has a higher water requirement and lower water-use efficiency than SO (Ponton *et al*., 2001, 2002). Furthermore, the availability of non-model plant species genomes, such as PO (Plomion *et al*., 2016), now allows an in depth analysis of the molecular mechanisms underlying stress responses in oaks including the (epi)genetic architecture of their ecological preferences (Estravis-Barcala *et al*., 2020). However most studies are so far limited to the analysis of the transcriptomic response of trees to abiotic stresses s as for example in gymnosperms (Behringer *et al*., 2015; Du *et al*., 2018; Fox *et al*., 2018) and in various angiosperms, including oaks (Villar *et al*., 2011; Torre *et al*., 2014; Tang *et al*., 2015; Mun *et al*., 2017; Müller *et al*., 2017; Guerrero-Sánchez *et al*., 2021). In this context, Epigenetics has recently emerged as essential regulatory mechanisms for plant development and for the responses and adaptations of plants to their environment (Richards *et al*., 2010). Epigenetics encompasses the non-DNA sequence-based heritable information carried by chromatin. It includes the posttranslational modification of histones, DNA methylation and specific small RNAs. In plants, DNA methylation occurs at cytosines in symmetrical (CG, and CHG), and non-symmetrical CHH sequence context (where H is A, C, or T) (Zhang *et al*., 2018). Genomic DNA methylation is known to be an essential component of the plant stress responses and memories (Gallusci et al, 2022; Zhang *et al*., 2018). Methylated cytosines are highly abundant in centromeres, and pericentromeric regions, which are enriched in transposable elements (TE). By contrast, methylation levels are low in euchromatic regions, characterized by higher gene densities (Niederhuth *et al*., 2016). It is generally accepted that DNA methylation plays an essential function in TE silencing but is also important for gene expression control (He et al., 2022).

So far, epigenetic studies have demonstrated that DNA methylation is highly dynamic when annual plants are subjected to abiotic stresses. In addition, mutants affected in *de novo* and/or maintenance DNA methylation present an altered response and adaptation to such stresses (Zhang *et al*., 2018; Kumar & Mohapatra, 2021). In trees, studies using poplar as a model system, demonstrated a general increase in DNA methylation during drought stress associated with a remodeling in DNA methylation patterns at both genes and TE, potentially leading to changes in the expression of genes involved in hormonal pathways (Sow *et al*., 2021, Liang *et al*., 2014; Lafon-Placette *et al*., 2018). Finally, the recent demonstration that *DDM1* knockdown results in genomic hypomethylation, and major changes in gene expression relative to wild type plants under DS, highlights the fundamental role of DNA methylation homeostasis in poplar trees facing DS (Sow *et al*., 2021).

At present, only two studies have investigated the epigenetic response of oak species to DS (Platt *et al*., 2015; Gugger *et al*., 2016). Both studies reported either Cpg polymorphisms or differentially methylated regions (DMRs) involved in local adaptation and potentially to drought resistance in *Quercus Lobata*.

However, there is no integrated analysis of the behavior of oak and more globally of forest trees, under DS. Here we present for the first time an integrated study, extending from the physiological characterization of oak plants and analysis of mRNA and small-RNA populations, to the distribution of methylated cytosines in PO and SO leaves. We used an experimental design, comparing the response of PO and SO grown under two contrasted water regimes: control, C (55 % of relative extractable water, REW) and drought stress, DS (35 % of REW). We observed major changes in mRNA and small-RNA populations associated with a remodeling of the methylome in both species in response to DS. The Main result highlights a strong species-specific epigenetic response to DS potentially in relation to the ecological preferences of PO and SO.

## Materials and methods

### Plant material and experimental design

The experiment was carried out using greenhouse grown PO and SO seedlings, subjected to a progressive soil drought stress (DS). Plants were grown in 2016 from seeds collected from pure French stands of each species, and transferred in spring 2017 to a greenhouse equipped with a robotic system for the automatic weighing and watering of plants (Bogeat-Triboulot *et al*., 2019). Leaves were harvested during a progressive decrease in soil water content: the first harvest date corresponds to a relative extractable water (REW) of 55%, a value well above the value of 40% below which forest trees reduce canopy transpiration when DS is developing (Granier *et al*., 2000). The second harvest date corresponds to a REW of 35%, which is below this 40% threshold (Fig. 1). Additional details concerning experimental design and monitoring of growth and gas exchanges are provided in supplementary information Methods S1 and S2. For nucleic acids (DNA, RNA and small RNAs) extraction, leaf blades were hand dissected to remove primary and secondary veins and immediately frozen in liquid nitrogen, and stored at -80°C until use.

**Figure 1:**
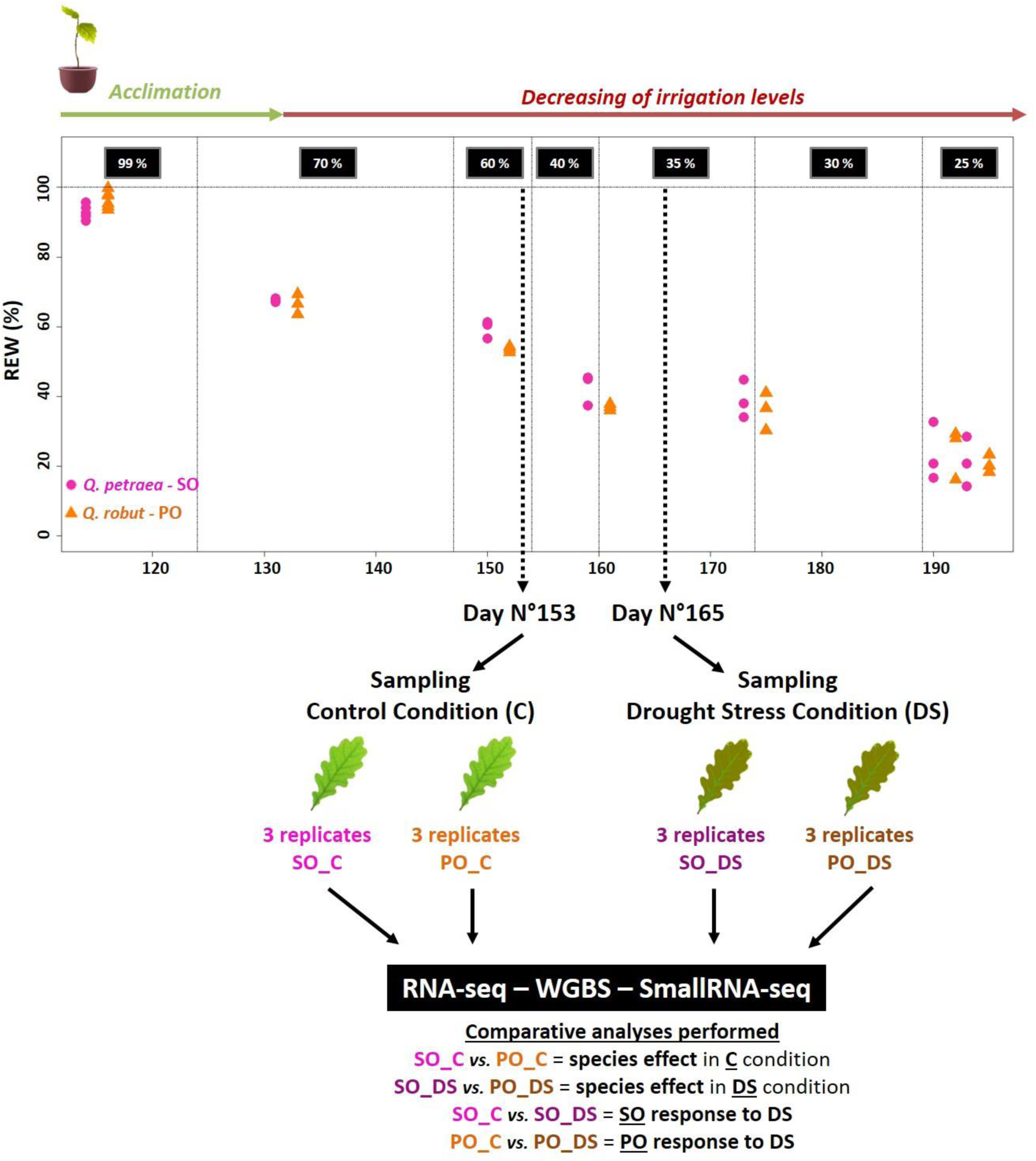
Characterization of the experimental design. The experiment took place over a six-month period, from April to August 2017 in a temperature controlledgreenhouse equipped with a robotic system for the automatic weighing and watering of plants (Bogeat-Triboulot et al., 2019). After watering to field capacity, a progressive drought stress was applied at day 132. Samples for molecular analysis (RNA-seq, SmallRNA-seq, WGBS) were collected from three plants of eachspecies on day 153 at a mean REW of 55 % (control conditions, C) and the second harvest on day 165 at a mean REW of 35% (drought stress, DS). For more details see Methods S1.

### Libraries construction and sequencing

RNA, small RNAs and genomic DNA were extracted using 100 mg powder with the mirVana RNA isolation kit (Thermo Fisher Scientific, USA) and the DNeasy plant minikit (Qiagen, Hilden, Germany), respectively according to manufacturer’s instructions (see Methods S3).

The 12 RNA and small-RNA (smRNA) libraries (2 species * 2 treatments *3 biological replicates) were generated and sequenced following standard procedure (Methods S3). Similarly, 12 libraries were generated for WGBS and sequenced as previously described (Daviaud *et al*., 2018).

### Transcriptome characterization

The sequence alignment files were generated with TMAP (Torrent Mapping Alignment Program, a dedicated mapper for ion torrent reads) (Caboche *et al*., 2014). Raw gene counts were generated with the featureCount function of the Rsubread package (Liao *et al*., 2014), which assigned the mapped sequencing reads to the gene models described in the PO genome (Plomion *et al*., 2016). The DESeq2 package (Love *et al*., 2014) was then used to identify differentially expressed genes. We investigated the effect of treatment and species in Wald tests (Love *et al*., 2014). Differentially expressed genes (DEGs) were identified with a false discovery rate (FDR)-adjusted p-value threshold of 0.05, on the basis of an absolute fold-change in expression > 1.5. Four gene sets were generated: (1) set#1, DEGs between control (C) and drought-stressed (DS) SO plants; (2) set#2, DEGs between C and DS PO plants; (3) set#3, DEGs between SO and PO plants grown in DS conditions; and (4) set#4, DEGs between SO and PO plants grown in C conditions.

### Small RNA-seq data processing

We used the same procedure described in (Rubio *et al*., 2022) for the quality treatment of the data. Remaining reads were mapped onto the *Q. robur* genome with Bowtie2. ShortStack (version v3.8.5) was used to identify and quantify small-RNA clusters with default settings (Axtell, 2013). Small-RNA clusters are uninterrupted linear genomic regions with a minimum sequencing depth of 20 reads. As clusters could be composed of a mixture of small RNAs of different sizes, a dicer call score was established: clusters for which at least 80% of reads were 20 to 24 nt long were considered to be dicer-derived, whereas all others were annotated as not dicer-derived and excluded from the analyses. ShortStack was also used to annotate miRNA loci according to a strict set of structural and expression-based criteria (Axtell, 2013).

### Whole-genome bisulfite sequencing (WGBS) analysis

After trimming (TrimGalore!, version v0.4.5), Bismark (version v0.20.0) (Krueger & Andrews, 2011) was used to calculate the bisulfite conversion rate using the unmethylated oak chloroplast genome. Alignment of cleaned reads with the PO reference genome was also performed with Bismark with a maximum of six mismatches. Reads for which multiple alignments were obtained were discarded as were PCR duplicates. The methylation state of each cytosine residue was calculated in each of the CG, CHG and CHH contexts. We used the DSS package (version 2.39.0) from R (Feng & Wu, 2019) to identify differentially methylated regions (DMRs) in a Wald test procedure, accounting for both biological variation between replicates and sequencing depths with standard parameters. For the definition of hypo- or hyperDMRs, we applied a cutoff for the difference in methylation ratio at least of 10% for CHH, and 25% for CHG and CG (Sow *et al*., 2021).

### Genomic location of DMRs and smRNA clusters

We used BEDTools (version v2.27.1) software and gff files: ‘gene body’ (defined in this study as exons + introns) and transposable elements (TE) from the PO reference genome. For both DMRs and smRNA clusters, BEDTools was used to identify the overlaps with gene bodies, 2 kb promoter regions and/or TEs, as described (Quinlan & Hall, 2010).

### Gene ontology enrichment analysis

We performed gene ontology (GO) enrichment analysis with the topGO package (Alexa, 2006). We performed Fisher’s exact tests to validate enrichment for each GO term (adjusted p-value < 0.05).

### Identification of overlaps between the transcriptome, methylome and small RNAs

We used BEDTools to identify overlaps between the transcriptome, methylome and small-RNA clusters. We performed three different analyses. The first aimed to identify the overlap between DMRs and DEGs, to highlight transcriptomic variation linked to methylation levels. The second aimed to identify overlaps between DMRs and smRNA clusters of 24 nt. We focused our analysis on the 24 nt smRNA clusters because these clusters are known to be involved in the methylation of the DNA regions they target. In the third analysis, we used the overlap obtained in the second analysis to test the overlap with DEGs, to highlight possible relationships between DEGs, 24 nt smRNA clusters and methylation levels.

### Fst

We investigated the possibility that the differential methylation between the two species arose through adaptive genetic differentiation, by investigating the overlap between loci differentially methylated between PO and SO independently of the treatment applied, and loci with high levels of genetic differentiation. The method used was described in a previous study (Le Provost *et al*., 2022).

## Results

### PO and SO respond differently to moderate drought stress

Drought stress (DS) was controlled to achieve drought levels close to the targeted values of Soil Relative Extractable Water 70, 60, 40, 35, 30 and 25% (Fig. S1a). During the humid control phase of the experiment (REW > 40%), PO had slightly higher CO2 assimilation rate (A), stomatal conductance for water vapor values (gs) and a lower water use efficiency (WUE) than SO. These differences became significant for gs at day 146 (Fig. S2). At 35% REW, just before the second harvest date (165 days), SO had a significantly higher gs, whereas after the harvest date, PO had a significantly higher A (Methods 2 and Fig S2). These data are consistent with the establishment of species specific responses when REW decreased from 55 % to 35%.

### The transcriptomic response to DS is essentially species-specific

The analysis of the leaf RNA population of PO and SO in C (control) and DS conditions were used to investigate eventual differences in the molecular responses of PO and SO to DS (Table S1). Principal component analysis (PCA), showed that all C samples except PO_C1 are grouped, suggesting low levels of variability between biological replicates, regardless of species (Fig. S3). In DS conditions, all PO and SO replicates, except PO_S3 and SO_S2, were clustered in two distinct groups along the PC1 axis. The DS_SO samples were barely separated from the PO and SO samples in C conditions along the PC2 axis. By contrast, DS_PO and C_PO samples are separated along the PC1 axis (Fig. S3), although scattered, suggesting a greater variability in the response to stress among PO genotypes.

We first determined the molecular response to DS within each of the two species. SO_DEGs correspond to differentially expressed genes (DEGs) in response to stress in SO, and PO_DEGs to those in PO species (Table S2). We identified 1845 DEGs in total, of which 1141 were PO_DEGs and 771 were SO_DEGs (Fig. 2a). Only a limited number of DEGs were common to SO_DEGs (27 %) and PO_DEGs (18 %). A majority of DEGs (55%) were upregulated in response to DS in SO species, whereas the converse was observed in PO, with 58% of downregulated DEGs.

**Figure 2:**
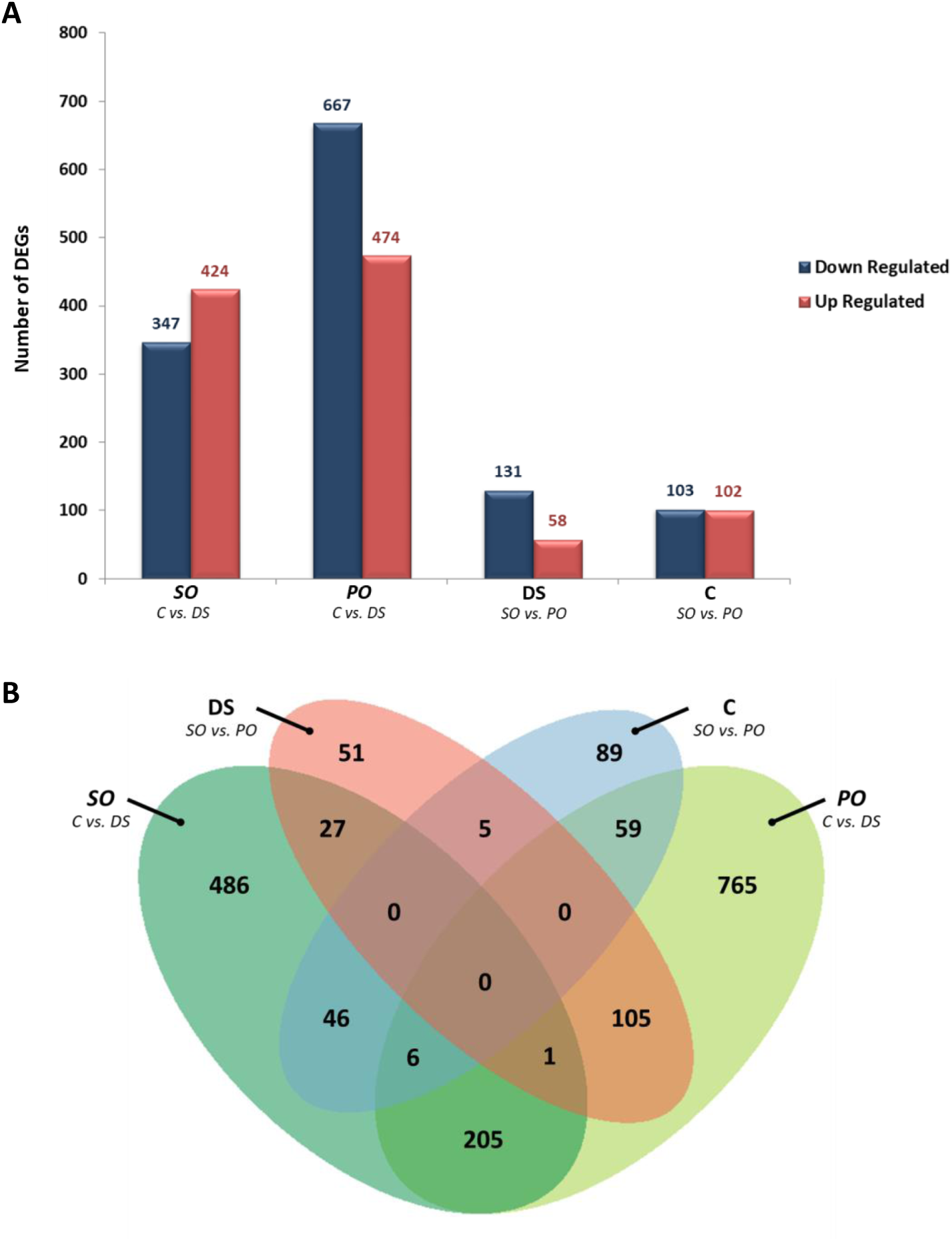
Summary of the differential gene expression analysis. Analysis were performed to identified DEGs related to response to DS in SO and PO. DEGs found comparing SO to PO in C and DS conditions were also reported**. (A)** DEGs identified in the four comparisons distinguishing down- (in blue) and up-regulated (in red) genes**. (B)** Venn diagram of the overlaps of all the DEGs of the four comparative analyses.

Gene ontology (GO) showed that SO_DEGs were enriched in genes involved in the ‘defense response’ (GO:0006852), ‘zinc ion transmembrane transport’ (GO:0071577) and ‘response to auxin’ (GO:0009733), whereas PO DEGs were mostly enriched in genes related to ‘metabolic process’ (GO:0008152), ‘protein phosphorylation’ (GO:0006468) and ‘oxidation-reduction process’ (GO:0055114) (Table S3-S4). Taken together, these results suggest that the molecular response to DS is mostly species-specific and reflects the difference in sensitivity to DS of these two species.

We then analyzed differences between species in each of the growing conditions. The C_DEGs correspond to genes differentially expressed between SO and PO in C conditions and DS_DEGs in DS conditions.

The total number of C_DEGs (205) and DS_DEGs (189) was similar but down-regulated DEGs accounted for 70% of the DS_DEGs whereas C_DEGS were equally distributed between up and down DEGs, consistent with a different trajectories of gene expression reprogramming in SO and PO under DS. (Fig. 2a). Most interspecific DEGs were condition-specific, with only 5 common DEGs identified (Fig. 2b) showing an extensive remodeling of gene expression profiles between the two conditions in PO and SO. Both the C and DS_DEGs were enriched in genes relating to ‘cell wall macromolecule catabolic process’ (GO:0016998) and ‘chitin catabolic process’ (GO:0006032’) (Table S3-S4).

### DNA methylation landscapes of SO and PO following DS

We used WGBS to analyze the methylation patterns in SO and PO under C and DS conditions (see methods). For all samples, the cytosine conversion rate after bisulfite treatment was over 99% (Table S5). The cytosine methylation levels were 57%, 36% and 5% in the CG, CHG and CHH contexts respectively, with no significant difference between species and/or conditions (Fig. S4). Methylation followed a bimodal distribution for CG and CHG whereas CHH methylation was weak (Fig. S5). We analyzed mean methylation patterns for genes and transposons (TEs) in each of the three C contexts. Overall, no significant differences were observed except for methylation levels in the gene body and TE in the CHH context (Fig. S6). In PCA, PO and SO were separated along the PC1 axis for the CG and CHG contexts, but not for CHH (Fig. S7) with a similar trend observed in the clustering analysis (Fig. S8). This suggests that within each species, methylome dynamics under DS conditions were driven in large part in an individual-dependent manner.

### Differentially methylated regions show species-specific responses to DS

We investigated the methylation dynamics of SO and PO to DS by analyzing DMRs (Differentially methylated regions) between the C and DS conditions in each species (Fig. 3). Four thousand eight hundred DMRs (3232 hypo- and 1568 hypermethylated) were identified in SO, and 4443 in PO (2464 hypo- and 1979 hypermethylated). Regardless of the species, DMRs in the CHH context were the most abundant, representing 67 % and 58 % of the total DMRs in SO and PO, respectively; Fig. 3). This suggests a higher dynamic of CHH methylation than in the two other contexts; an observation also confirmed when DMCs are analyzed..

**Figure 3:**
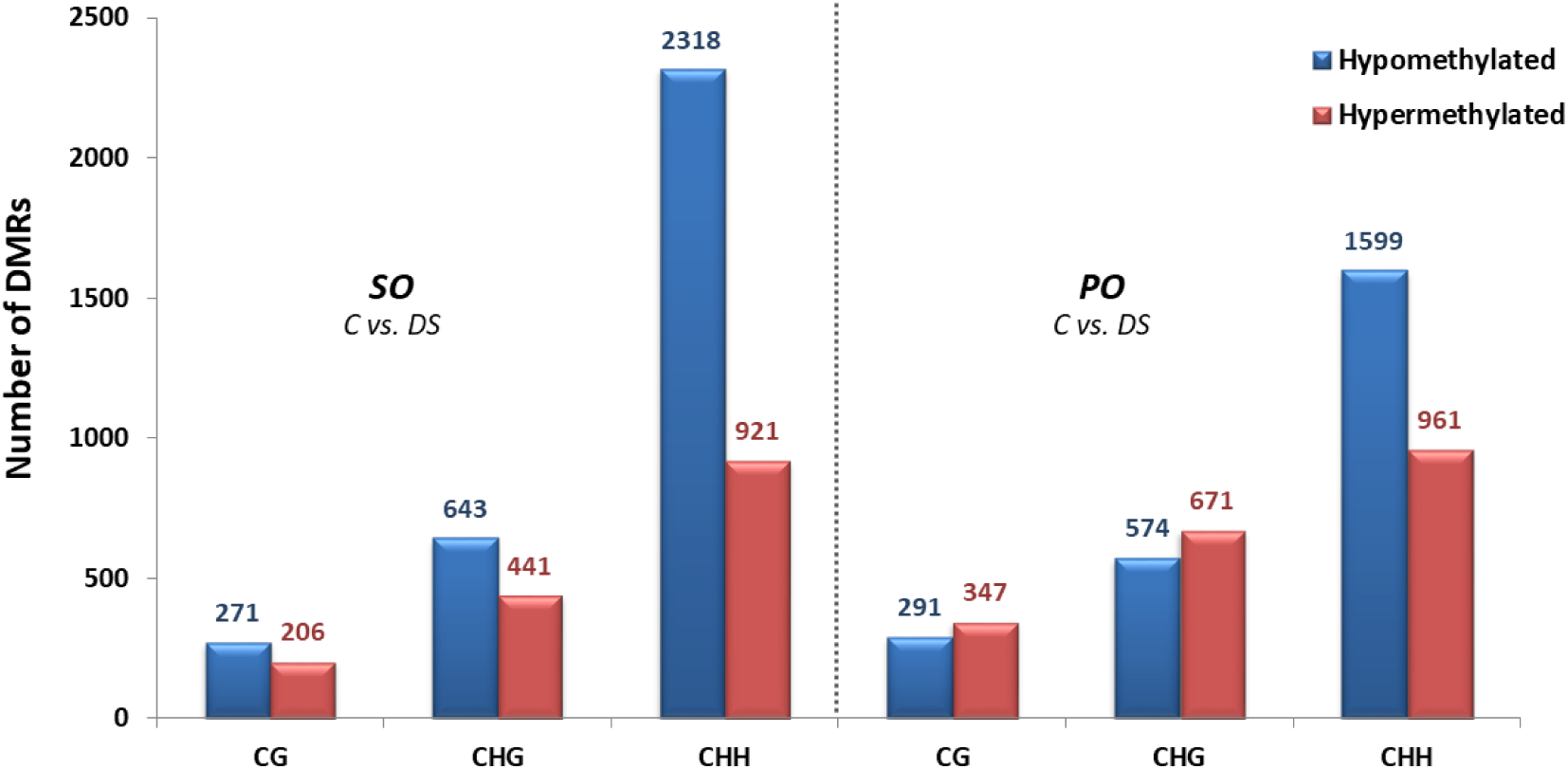
Differentially methylated regions between C and DS conditions of each species. DMRs have been differentiated in hypomethylated (in blue) and hypermethylated categories.

The SO-hypo-DMRs are slightly more abundant than hyper-DMRs in the symmetrical sequence contexts, but are by far the most abundant in the CHH sequence context where they represent 71.5% of theDMRs.In PO, hyper-DMRs represent 53% and 54 % of DMRs in the CG and CHG contexts, whereas a majority of DMRs (63.6%) are hypomethylated in the CHH context (Fig. 3). In both species, most DMRs were located in TEs (70% for SO and 75% for PO). However a significant proportion of DMRs is also found in 2 kb promoter regions (∼11.5% for SO and 10.1% for PO) and in gene bodies (∼ 7.1% for SO and 4.6% for PO) (Table S6). Taken together these data show that there are no major differences between both species, except for a higher number of DMRs in the CHH context in SO than in PO. However, analyzing the overlap between the SO and PO_DMRs in each of the three C sequence contexts (Fig. 4a) indicated a very limited number of DMRs common to the two species (2, 8 and 143 in the CG, CHG and CHH contexts, respectively). This is consistent with a species-specific epigenetic response to DS.

**Figure 4:**
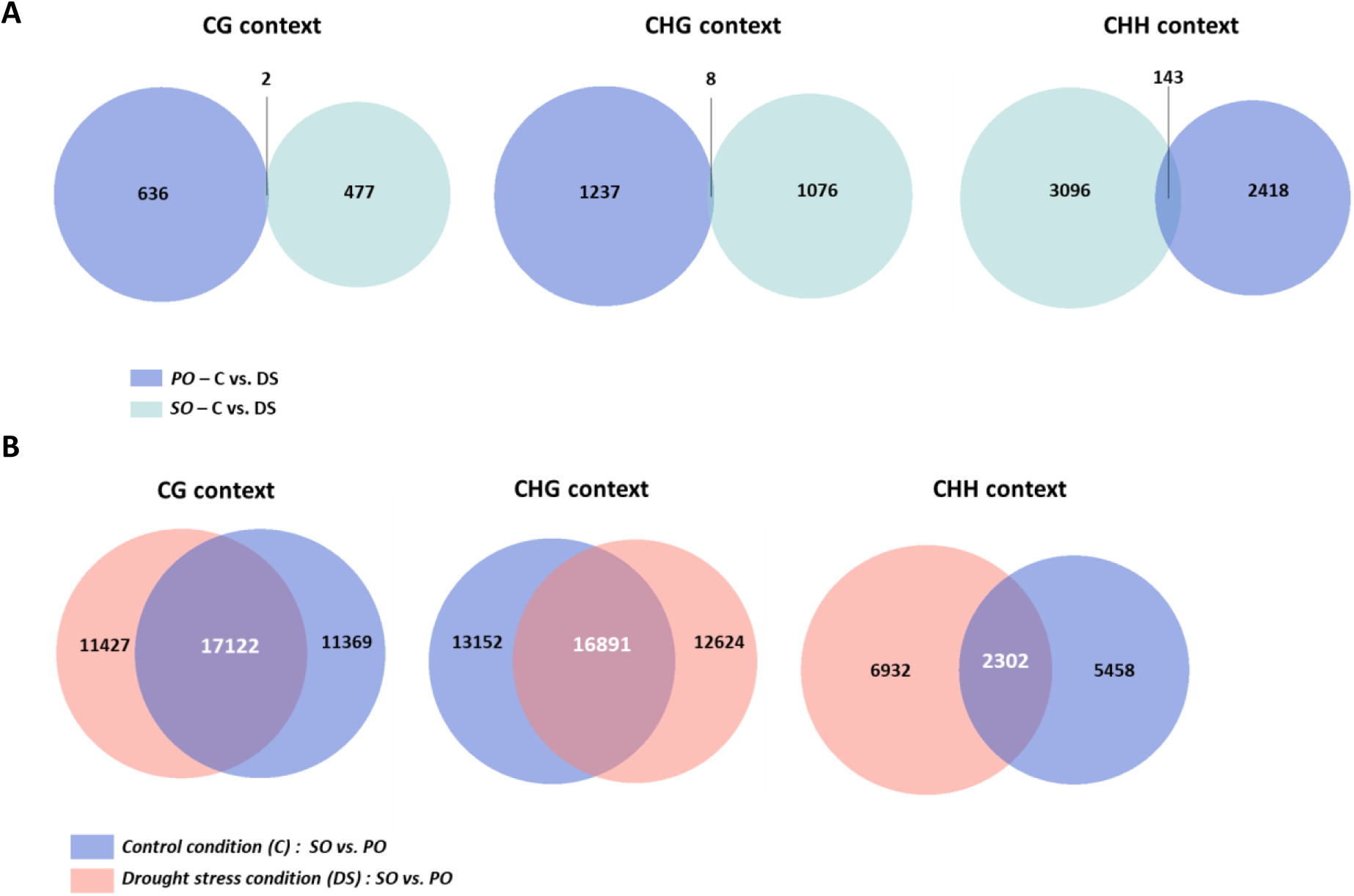
Venn diagrams of the overlaps of (A) DMRs obtained from SO (in green) and PO (in blue) after comparions between treatments and (B) by comparing SO and PO in control (in blue) and drought stress (in orange) conditions. The search for overlaps was performed in CG, CHG and CHH context.

### Many interspecific DMRs are conditions independant

We identified DMRs between SO and PO in C (C-DMRs) and in DS (DS-DMRs) conditions. The total number of C_DMRs and DS_DMRs was similar with a balanced distribution between hypo- and hyperDMRs (Fig. 5). The CHG and CHH C_DMRs and DS_DMRs were mostly hypomethylated (51%) and hypermethylated (52%), respectively, in SO, with no inversion between conditions. By contrast, hypermethylated CG C_DMRs were more abundant in SO (60%) than in PO (41%), but the opposite pattern was observed in DS conditions, with 57% hypomethylated CG DS_DMRs in SO and 43% in PO (Fig 5). To better characterize the inversion of the ratio between hypo- and hypermethylated CG DMRs, their genomic locations were analyzed (Table S7). For annotated CG DMRs, similar genomic locations were identified for C- and DS-DMRs, with most of the DMRs located in TEs (∼ 69.1%), in 2 kb promoter regions (∼ 10.1%) and in gene bodies (∼ 10.7%) (Table S7).

**Figure 5:**
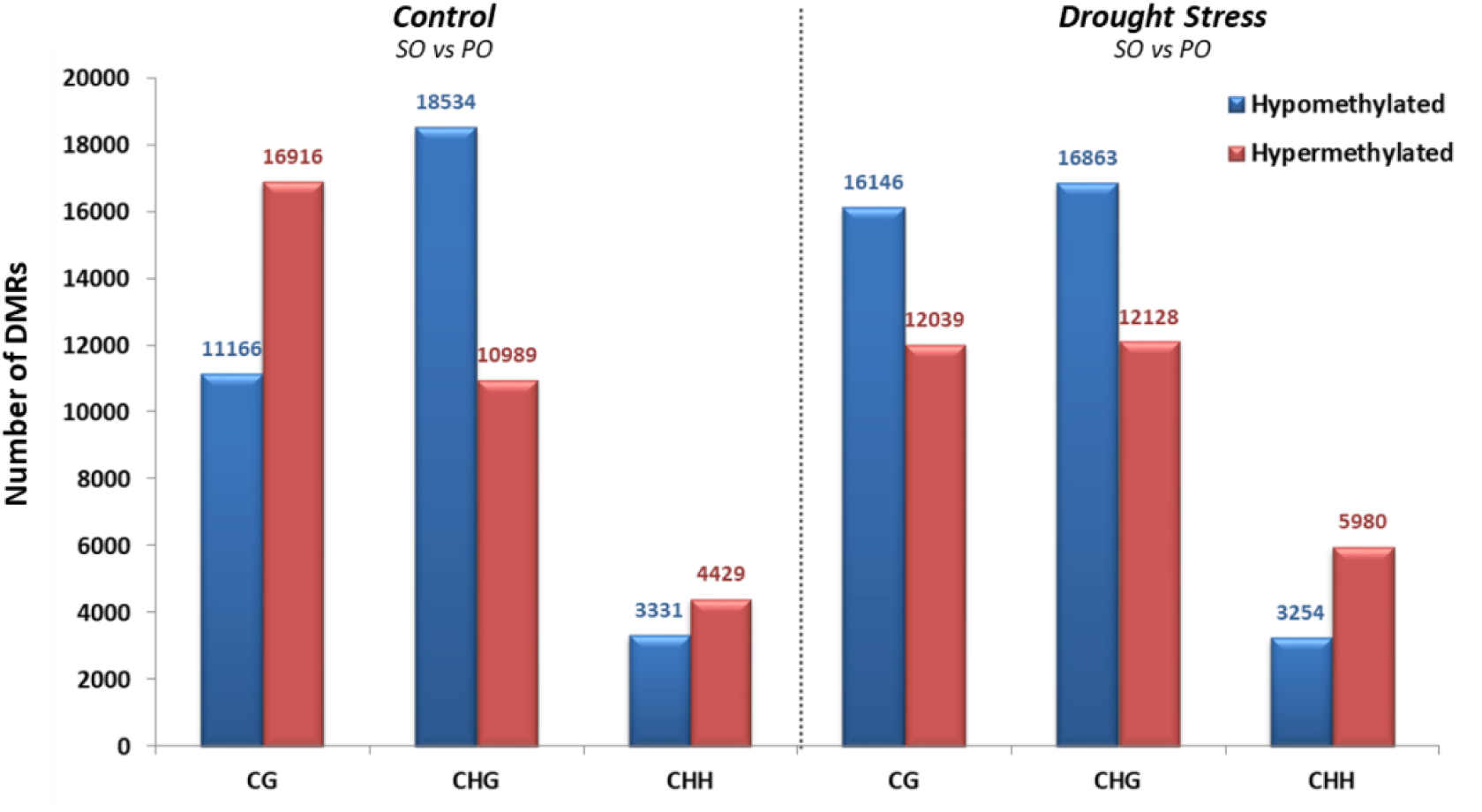
Differentially methylated regions between SO and PO in control and drought stress conditions. DMRs have been differentiated in hypomethylated (in blue) and hypermethylated categories.

We then investigated whether C-DMRs were maintained in DS conditions as they may reflect fundamental epigenetic differences between PO and SO regardless of their growing environment. The overlap between C- and DS-DMRs accounts for 60% and 30% of the DMRs identified in the CG/CHG and CHH contexts, respectively (Fig. 4b), of which 97 (0.3%) displayed inversions of methylation status according to growing conditions (i.e. 54 and 43 became hyper- and hypomethylated in DS conditions, respectively) (Table S8)..

The 36315 DMRs common to both conditions were located for most of them in TE (51%), in gene bodies (9.7 %) and in 2 kb promoter regions (5.5%) (Table S9). GO enrichment analyses of DMRs annotated in gene bodies and in 2 kb promoter regions (Table S11) revealed an enrichment in terms relating to ‘protein phosphorylation’ (GO :0006468), ‘signal transduction’ (GO:0007165), ‘response to auxin’ (GO:0009733) and ‘carbohydrate metabolic process’ (GO:0005975).

Because these stable methylation marks are species specific, we investigated their genomic regions. A significant enrichment of genes including highly differentiated genetic markers between populations of the two oak species was found among the genes presenting the stable DMRs. (Table S10).

### Correlation between DNA methylation and gene expression

We assessed the relationship between gene expression and DNA methylation by analyzing the genomic colocalization between DEGs and DMRs (DMEGs for differentially methylated and expressed genes) for the four comparisons: SO_C/DS, PO_C/DS, DS_SO/PO and C_SO/PO. We determined the coordinates of the gene body (DEGs_gb) and the 2 kb promoter region (DEG_2kbprom) for each DEG and performed overlap with DMRs. In total, 124 DMEGs_Gb and 156 DMEGs_Prom were identified (Table S12 and S13).

Boxplots generated between methylation status and the log fold-change in expression of the DEGs for both the DMEGs_Gb (Fig. 6a) and the DMEGs_Prom (Fig. 6b) highlighted significant differences in DS conditions. Among the DMEGs_Gb and the DMEGs_Prom, 17/18 and 18/21 respectively, were hypermethylated and repressed in SO versus PO. They included genes encoding tyrosine kinases, chitinases and F-box-associated domain proteins and cytoskeletal regulator Flightless-I proteins (Table S13). In addition, 15/27 DMEGs_Gb and 14/21 DMEGs_Prom which include genes encoding cytoskeletal regulator Flightless-I proteins and NAD-dependent malate dehydrogenases, were hypomethylated and overexpressed in SO versus PO (Table S13).

**Figure 6:**
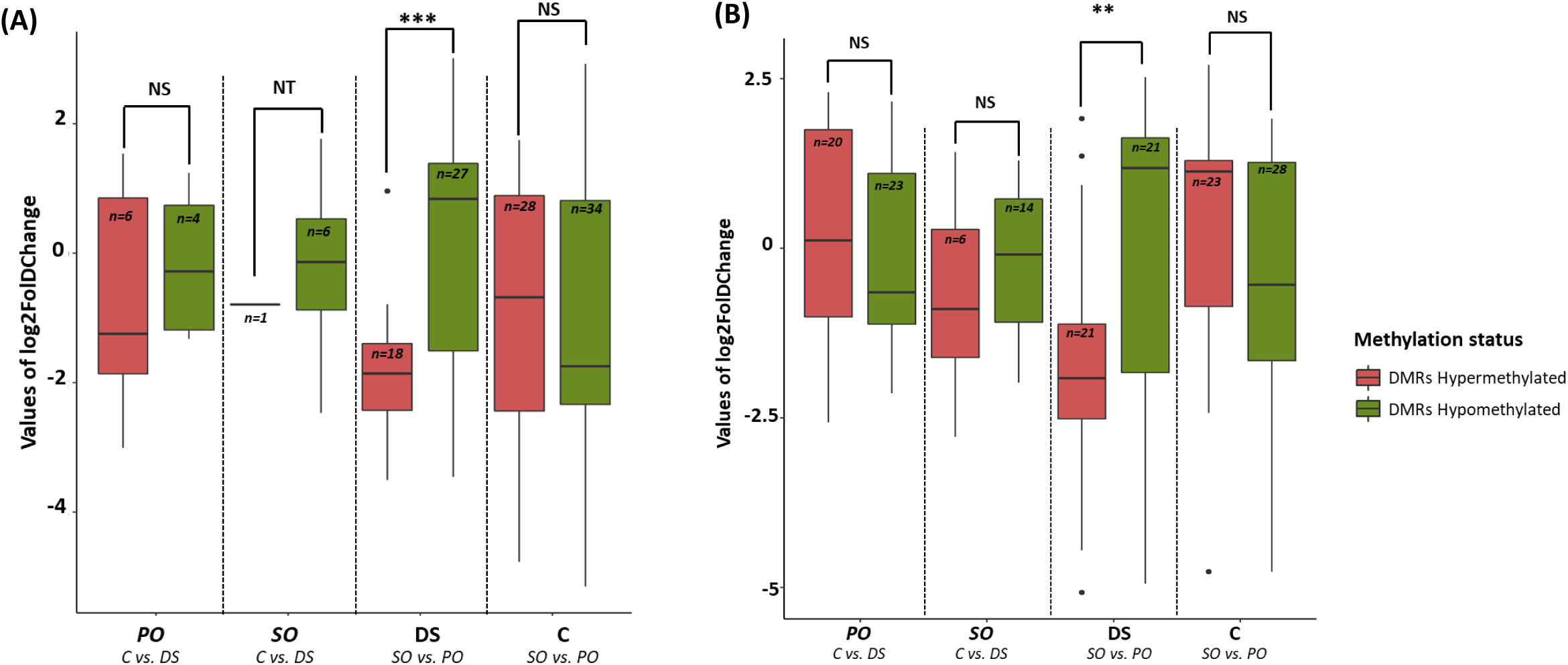
Box plot of differential expression levels of hypo- and hypermethylated DMRs identified in the different comparative analyzes. The differential expression levels were represneted in the y-axis (Values of log2FoldChange). The hyper- and hypomethylated DMRs are presented by red and green box plots respectively.The significative statistical differences of the differential expression levels between hyper- and hypomethylated DMRs were performed using Wilcoxon-Mann-Whitney tests.Significant codes : 0 ‘***’ 0.001 ‘**’ 0.01 ‘*’ 0.05 ‘.’ 0.1 NS : No significant, NT : No tested.

### Small-RNA populations differs between SO and PO in C and DS conditions

Small RNA populations were characterized for SO and PO in C and DS conditions (Table S14). The siRNA and miRNA clusters were determined by merging the data from all replicates in each species and treatment, to generate four smRNA cluster populations (2 species by 2 conditions). We took into account the difference in read numbers between samples by considering only clusters with (1) a minimum read number of three in each replicate and (2) a coefficient of variation of no more than 50% (i.e. defined as the standard deviation for the three replicates divided by the mean of the three replicates). In total 37978, 34660, 39966 and 52925 smRNA clusters were identified for SO_C, SO_DS, PO_C and PO_DS, respectively (Table S15).

As described by Axtell (2013), miRNA and siRNA clusters were differentiated with miRNA clusters encompassing mostly reads of 21 nt (57% - 64%) whereas siRNA clusters contain mostly reads of 24 nt (96% - 98%) (Fig. S9) as reported in grapevine (Rubio *et al*., 2022), *Arabidopsis*, rice, tomato and maize (Axtell, 2013). For half of the clusters matching with PO reference genome annotation (Table S16), most of them were annotated as TEs (∼ 68%) followed by promoters (∼14.3%) and gene bodies (∼ 5.5%) with a distribution similar in both species and conditions (Fig. S10).

To investigate the effects of conditions on smRNA populations, we considered only qualitative variations (i.e. the presence or absence of smRNA clusters). A large proportion of the clusters identified in SO plants were common to both the C (66%) and DS (72%) conditions (Fig. 7). Among them, only 1196 clusters (∼ 4.8% of the clusters common to C and DS conditions) had a FC ratio of at least 2, suggesting a limited impact of DS on smRNA populations in SO. By contrast, DS resulted in a large change in the PO smRNA population, as demonstrated by the small proportion of clusters common to both sets of conditions (57% in C conditions and 43% in DS conditions), consistent with the greater sensitivity to DS of this species. Interspecific comparisons, regardless of the conditions considered, highlighted the large proportion of smRNAs that were species-species specific. In control conditions, 88.5% of smRNA clusters were specific to SO, and 91.4% were specific to PO. A similar trend was observed in DS conditions, with 71.3% to 91% of clusters specific to SO and PO, respectively (Fig. 7).

**Figure 7:**
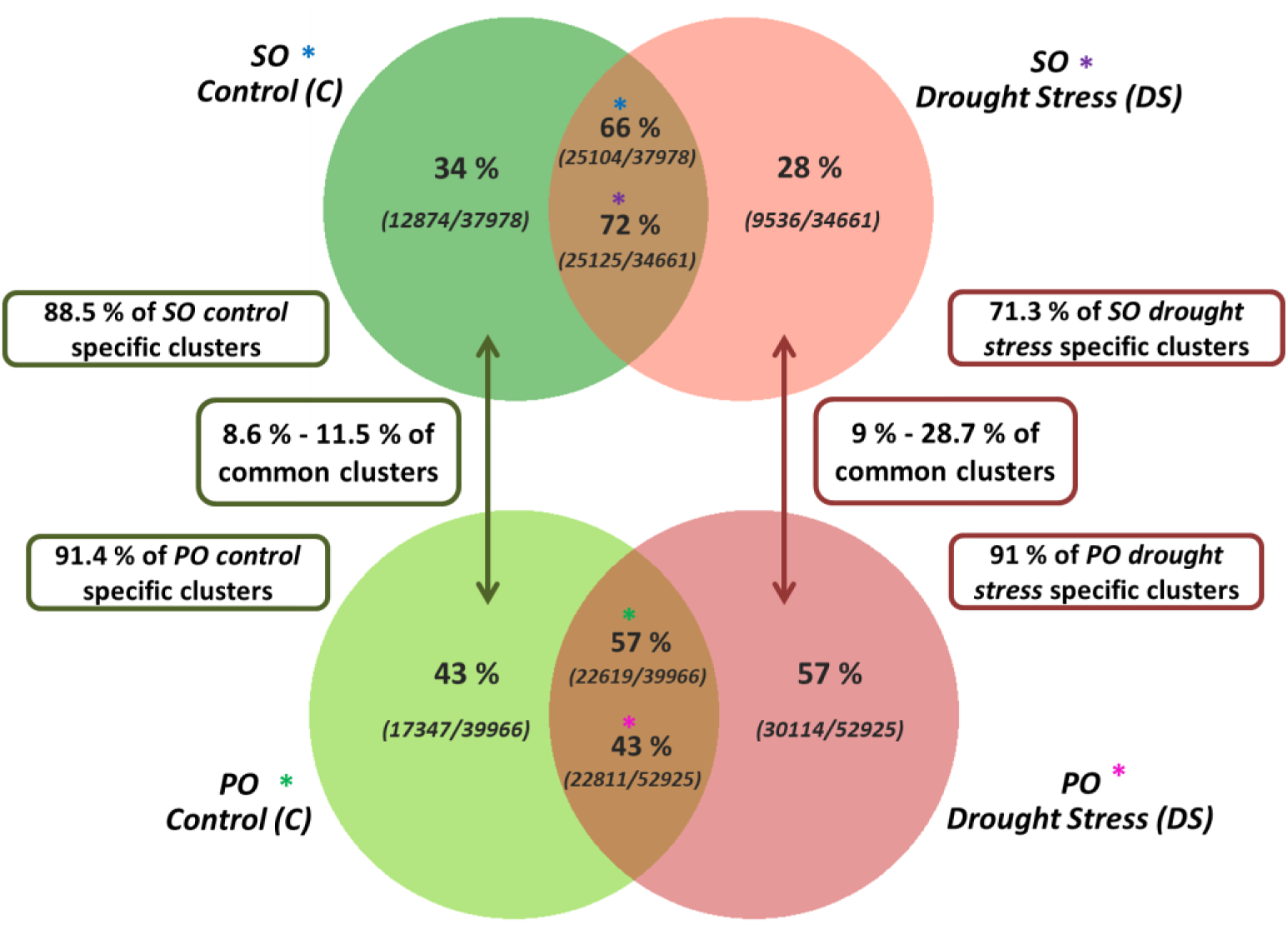
Comparions of small RNA clusters in control and drought stress in SO and PO species. The top Venn diagram corresponds to the comparison of the clusters (absence/presence) betweencontrol and drought stress in SO. The botton diagram corresponds to the comparison of the clustersbetween in control and drought stress in PO. The clusters identified in common between before and after water stress conditions are referenced by blue, violet, green and pink stars for SO_C, SO_DS, PO_C, PO_DS, respectively. The results of the overlap search between the specific clusters of each species in C condition are presented in the green boxes. The results of the overlapsearch between the specific clusters of each species in DS condition are presented in the red boxes.

GO terms enrichment analysis were performed on common (Table S17) and specific smRNA clusters (Tables S18 and S19). The results are consistent with the previous observation that PO and SO deployed different molecular mechanisms in response to DS.

### Correlation between 24 nt smRNAs and DNA methylation

We assessed the potential association between 24-nt smRNAs and DNA methylation by looking for overlaps between 24-nt smRNA clusters and DMRs.. We applied a three-step bottleneck approach to identify the most relevant overlaps, defined here as those with a 24-nt smRNA cluster specific to a single condition in a selected comparison (eg: PO-C specific cluster in the comparison between PO_C and PO_DS) and associated with the hypermethylation in the same condition of DMRs identified in the same comparison (Fig. S11a). In total, 8443 overlaps were identified (Fig. S11b). In both species, most of them (70% to 94%) were associated with CHH-DMRs in response to DS. In contrast, overlaps were evenly distributed between the three contexts when the two species were compared in C and DS conditions respectively The overlaps identified in PO_C and PO_DS were five and three times more abundant than those in SO_C and SO_DS, respectively (Fig. S11b).

The 8443 regions overlapping with both DMRs and small-RNAs clusters were also analyzed for colocalization with DEGs, based on the coordinates of the gene body and the 2 kb promoter region, respectively. We found that 26 hypermethylated DMRs overlapped with both a smRNA cluster and a downregulated DEG among which 14 overlapped with gene bodies and 12 with promoter regions (Table 1).

**Table 1:**
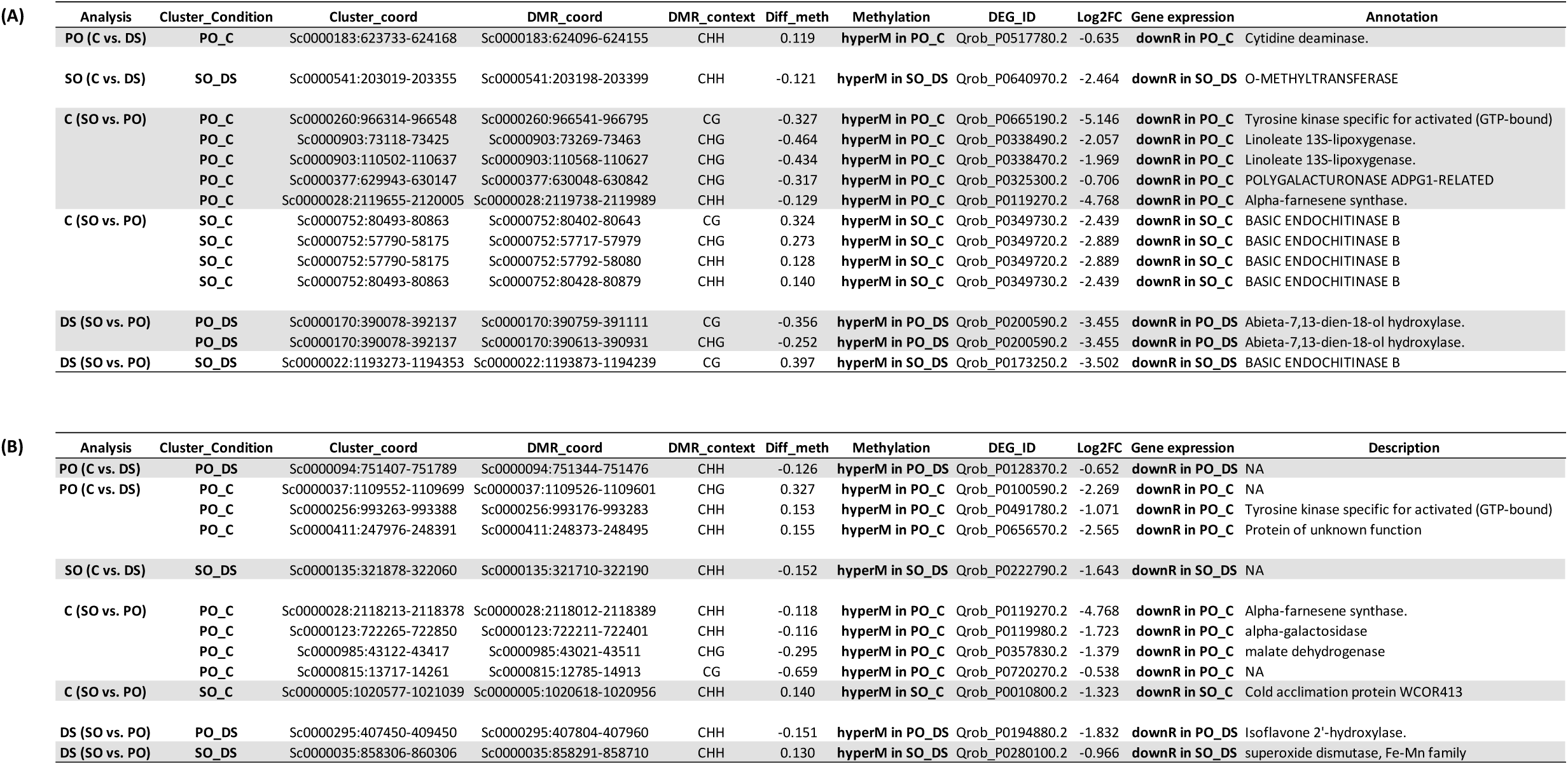
Hypermethylated regions associated with the presence of a 24-nt smRNA cluster co- localizing withan under-expressed DEG at the gene body level (A) and at the 2 kb promoter region (B).

| Analysis | Cluster_Condition | Cluster_coord | DMR_coord | DMR_context | Diff_meth | Methylation | DEG_ID | Log2FC | Gene expression | Annotation |
| --- | --- | --- | --- | --- | --- | --- | --- | --- | --- | --- |
| PO (C vs. DS) | PO_C | Sc0000183:623733-624168 | Sc0000183:624096-624155 | CHH | 0.119 | hyperM in PO_C | Qrob_P0517780.2 | -0.635 | downR in PO_C | Cytidine deaminase. |
| SO (C vs. DS) | SO_DS | Sc0000541:203019-203355 | Sc0000541:203198-203399 | CHH | -0.121 | hyperM in SO_DS | Qrob_P0640970.2 | -2.464 | downR in SO_DS | O-METHYLTRANSFERASE |
| C (SO vs. PO) | PO_C | Sc0000260:966314-966548 | Sc0000260:966541-966795 | CG | -0.327 | hyperM in PO_C | Qrob_P0665190.2 | -5.146 | downR in PO_C | Tyrosine kinase specific for activated (GTP-bound) |
|  | PO_C | Sc0000903:73118-73425 | Sc0000903:73269-73463 | CHG | -0.464 | hyperM in PO_C | Qrob_P0338490.2 | -2.057 | downR in PO_C | Linoleate 13S-lipoxygenase. |
|  | PO_C | Sc0000903:110502-110637 | Sc0000903:110568-110627 | CHG | -0.434 | hyperM in PO_C | Qrob_P0338470.2 | -1.969 | downR in PO_C | Linoleate 13S-lipoxygenase. |
|  | PO_C | Sc0000377:629943-630147 | Sc0000377:630048-630842 | CHG | -0.317 | hyperM in PO_C | Qrob_P0325300.2 | -0.706 | downR in PO_C | POLYGALACTURONASE ADPG1-RELATED |
|  | PO_C | Sc0000028:2119655-2120005 | Sc0000028:2119738-2119989 | CHH | -0.129 | hyperM in PO_C | Qrob_P0119270.2 | -4.768 | downR in PO_C | Alpha-farnesene synthase. |
| C (SO vs. PO) | SO_C | Sc0000752:80493-80863 | Sc0000752:80402-80643 | CG | 0.324 | hyperM in SO_C | Qrob_P0349730.2 | -2.439 | downR in SO_C | BASIC ENDOCHITINASE B |
|  | SO_C | Sc0000752:57790-58175 | Sc0000752:57717-57979 | CHG | 0.273 | hyperM in SO_C | Qrob_P0349720.2 | -2.889 | downR in SO_C | BASIC ENDOCHITINASE B |
|  | SO_C | Sc0000752:57790-58175 | Sc0000752:57792-58080 | CHH | 0.128 | hyperM in SO_C | Qrob_P0349720.2 | -2.889 | downR in SO_C | BASIC ENDOCHITINASE B |
|  | SO_C | Sc0000752:80493-80863 | Sc0000752:80428-80879 | CHH | 0.140 | hyperM in SO_C | Qrob_P0349730.2 | -2.439 | downR in SO_C | BASIC ENDOCHITINASE B |
| DS (SO vs. PO) | PO_DS | Sc0000170:390078-392137 | Sc0000170:390759-391111 | CG | -0.356 | hyperM in PO_DS | Qrob_P0200590.2 | -3.455 | downR in PO_DS | Abieta-7,13-dien-18-ol hydroxylase. |
|  | PO_DS | Sc0000170:390078-392137 | Sc0000170:390613-390931 | CHG | -0.252 | hyperM in PO_DS | Qrob_P0200590.2 | -3.455 | downR in PO_DS | Abieta-7,13-dien-18-ol hydroxylase. |
| DS (SO vs. PO) | SO_DS | Sc0000022:1193273-1194353 | Sc0000022:1193873-1194239 | CG | 0.397 | hyperM in SO_DS | Qrob_P0173250.2 | -3.502 | downR in SO_DS | BASIC ENDOCHITINASE B |

(B)
| Analysis | Cluster_Condition | Cluster_coord | DMR_coord | DMR_context | Diff_meth | Methylation | DEG_ID | Log2FC | Gene expression | Description |
| --- | --- | --- | --- | --- | --- | --- | --- | --- | --- | --- |
| PO (C vs. DS) | PO_DS | Sc0000094:751407-751789 | Sc0000094:751344-751476 | CHH | -0.126 | hyperM in PO_DS | Qrob_P0128370.2 | -0.652 | downR in PO_DS | NA |
| PO (C vs. DS) | PO_C | Sc0000037:1109552-1109699 | Sc0000037:1109526-1109601 | CHG | 0.327 | hyperM in PO_C | Qrob_P0100590.2 | -2.269 | downR in PO_C | NA |
|  | PO_C | Sc0000256:993263-993388 | Sc0000256:993176-993283 | CHH | 0.153 | hyperM in PO_C | Qrob_P0491780.2 | -1.071 | downR in PO_C | Tyrosine kinase specific for activated (GTP-bound) |
|  | PO_C | Sc0000411:247976-248391 | Sc0000411:248373-248495 | CHH | 0.155 | hyperM in PO_C | Qrob_P0656570.2 | -2.565 | downR in PO_C | Protein of unknown function |
| SO (C vs. DS) | SO_DS | Sc0000135:321878-322060 | Sc0000135:321710-322190 | CHH | -0.152 | hyperM in SO_DS | Qrob_P0222790.2 | -1.643 | downR in SO_DS | NA |
| C (SO vs. PO) | PO_C | Sc0000028:2118213-2118378 | Sc0000028:2118012-2118389 | CHH | -0.118 | hyperM in PO_C | Qrob_P0119270.2 | -4.768 | downR in PO_C | Alpha-farnesene synthase. |
|  | PO_C | Sc0000123:722265-722850 | Sc0000123:722211-722401 | CHH | -0.116 | hyperM in PO_C | Qrob_P0119980.2 | -1.723 | downR in PO_C | alpha-galactosidase |
|  | PO_C | Sc0000985:43122-43417 | Sc0000985:43021-43511 | CHG | -0.295 | hyperM in PO_C | Qrob_P0357830.2 | -1.379 | downR in PO_C | malate dehydrogenase |
|  | PO_C | Sc0000815:13717-14261 | Sc0000815:12785-14913 | CG | -0.659 | hyperM in PO_C | Qrob_P0720270.2 | -0.538 | downR in PO_C | NA |
| C (SO vs. PO) | SO_C | Sc0000005:1020577-1021039 | Sc0000005:1020618-1020956 | CHH | 0.140 | hyperM in SO_C | Qrob_P0010800.2 | -1.323 | downR in SO_C | Cold acclimation protein WCOR413 |
| DS (SO vs. PO) | PO_DS | Sc0000295:407450-409450 | Sc0000295:407804-407960 | CHH | -0.151 | hyperM in PO_DS | Qrob_P0194880.2 | -1.832 | downR in PO_DS | Isoflavone 2'-hydroxylase. |
| DS (SO vs. PO) | SO_DS | Sc0000035:858306-860306 | Sc0000035:858291-858710 | CHH | 0.130 | hyperM in SO_DS | Qrob_P0280100.2 | -0.966 | downR in SO_DS | superoxide dismutase, Fe-Mn family |

## Discussion

Integrative approaches are still in their infancy for non-model species, such as forest trees. The development of high-throughput sequencing technology now allows innovative (epi)genomic research on local adaptations of species, as genomic information on non-model species is accumulating <u>(Sork, 2017)</u> . Consequently, a new disciplinary field — ecological epigenomics — is now emerging, that aims at integrating epigenomics in the analysis of ecologically relevant phenotypic variations and at predicting evolutionary trajectories (Lamka *et al*.). In this context, the European white oak system, consisting principally of the two species PO and SO, constitutes a model of choice for analyzing the potential contribution of epigenomic differences to adaptation to contrasting environments.

We took advantage of this situation to investigate the specificity of drought responses in PO and SO by integrating for the the first time in oak species steady-state transcript levels, DNA methylation profiles and small-RNAs . Our data are consistent with a species-specific response to drought stress. In addition, the genomic colocalization of smRNAs, DMRs and DEGs in the two species may reflect the existence of important regions for forest tree adaptation to DS.

### Gene expression profiles are consistent with a higher sensitivity to DS in PO than in SO

Comparison of the transcriptomic response to DS in each species indicates that there are twice more upregulated DEGs in PO than in SO. Similar results were obtained comparing the response to DS of PO to two different well-known DS-tolerant oak species (*Q. ilex* and *Q. pubescens, (*<u>Madritsch *et al*., 2019</u>)), and the ecological differences between PO and other DS sensitive species, including S0, in their natural environment (Eaton *et al*., 2016).

In addition, the transcriptomic response to DS was in large part species-specific with 62% of DEGs exclusive to SO and 76% to PO, as were the physiological processes identified in GO term enrichment analysis. Indeed many processes are common between both species when common DEGs are analyzed.

However, many processes appear species specific. The most significant GO terms in PO are related to metabolic processes, protein phosphorylation and to the oxidative stress response. They are mostly driven by genes encoding proteins relating to oxidation mechanisms (laccases, cytochrome P450 and deacetoxy vindoline 4-hydroxylase) indicating that PO plants are subject to a strong oxidative stress during DS, as in model plants (Sharma *et al*., 2012; Pandian *et al*., 2020; Arcuri *et al*., 2020; Nadarajah, 2020). Many upregulated genes encode tyrosine kinases, as in arabidopsis and rice (Rodríguez & Canales, 2005; Allimuthu *et al*., 2020). By contrast, the analysis of the GO terms in SO, which correspond to a smaller number of genes than in PO, showed an enrichment in terms relating to “defense response”, “response to auxin” and “NADP biosynthetic process”. Taken together, these results suggest that the molecular mechanisms mobilized in response to DS differ between these two species and reflect their ecological differences, as reported for the contrasting DS ecotypes in *Q. lobata* (Gugger *et al*., 2017).

Comparison of PO and SO in C and DS conditions also indicates that their transcriptomic trajectory under DS is very different as only very few genes are common between C- and DS-DEGs (5 DEGs, Fig.2b). This is also demonstrated by the observation that, although some biological functions are mobilized in response to DS in both species (2 from the 4 and 7 GO identified in SO and PO, respectively), they are driven by different members of multigenic families (Table S3). In particular, genes encoding chitinases and endochitinases were found to be upregulated in PO regardless of growing conditions as described by Gugger et al. (2017) in *Q. lobata* in response to DS.

Finally, for C-DEGs, genes encoding cellulose synthases were found to be upregulated in SO relative to PO. Cellulose synthesis may be crucial for cell wall structure and the maintenance of cell turgor under low water potential, allowing continuous cell growth (Le Gall *et al*., 2015; Wang *et al*., 2016; Kesten *et al*., 2017; Ezquer *et al*., 2020).

### Analysis of DNA methylation demonstrates species-specific methylome dynamics under drought stress

Although DMRs identified following DS were present in similar numbers in the two species (4800 SO_DMRs and 4443 PO_DMRs, Fig 3), they present species-specific features. (1) the proportion of DMRs is each sequence context differs markedly in PO (14.3%, 28%, 57.7%; in the CG, CHG and CHH contexts respectively) and SO (9.9%, 22.5%, 67.5%; in the CG, CHG and CHH contexts respectively); this result highlights the difference in CHH methylation dynamics between SO and PO and suggests more dynamic changes in CHH methylation than in the two other contexts as previously described in mungbean (Zhao *et al*., 2022) and cotton (Lu *et al*., 2017). However this effect is more marked in SO than in PO ; (2) the proportion of hypo/hyper CHH DMR ratio differs between species (2.5 in SO and 1.7 in PO). This difference is due to a 1.5 fold enrichment in CHH hypermethylated DMRs in SO compared to PO under DS, whereas hypo DMRs are in similar numbers in both species. This observation suggests that the RdDM pathway is more active in SO than in PO under DS, (3) the ratio between hypo and hyper DMRs in the CG and CHG sequence contexts is opposite.Taken together these results suggest that white oaks present species specific DNA methylation dynamics under drought stress.

### Methylome variations and gene expression dynamics are positively correlated in drought stress for a set of candidate genes

We explored the relationship between methylation changes and transcriptional dynamics, by investigating the overlap between DMRs and DEGs. Our data showed that there is a significant correlation between DNA methylation level and gene expression for 32 DMEGs_Gb and 32 DMEGs_prom (Fig. 6 and Table S12). Among the 17 and 18 hypermethylated DMEGs_Gb and DMEGs_prom, overexpressed in PO two are of particular interest (i) a no apical meristem protein (NAM) and (ii) a transcription initiation factor, TFIID. The NAM proteins belong to the the largest plant-specific NAC TF family and are involved in many plant developmental processes and in (a)biotic stress responses (Singh & Laxmi, 2015; Tweneboah & Oh, 2017). They have been involved in DS responses in annuals (Puranik *et al*., 2012). The involvement of TFIID encoding genes in DS responses have been shown for *A. thaliana* (Gao et al., 2006), finger millet (Parvathi *et al*., 2019) and rice (Zhang *et al*., 2020). These genes could contribute to providing SO species a better tolerance to drought.

Focusing on the 15 and 14 hypomethylated DMEGs_Gb and DMEGs_prom, respectively overexpressed in SO, many encode cytoskeletal proteins. The plant cytoskeleton is essential for the maintenance of cell structure. It has three major structural components: microtubules, actin and intermediate filaments (Soda *et al*., 2016). Microtubules are a key sensor of stress responses in plants. In response to DS, they regulate plant stomatal morphology, cell wall construction and the accumulation of abscisic acid (Ma & Liu, 2019). These cell wall elements may also be key molecular players in the response of SO to DS, consistent with the results of our transcriptome analysis, which highlighted a role for genes relating to the cellulose synthase process.

We also identified a gene encoding a protein with a KDEL target peptide sequence, which prevents protein secretion from the endoplasmic reticulum (ER) and facilitates the return of the protein to the ER if accidentally exported (Yamamoto *et al*., 2003). Abiotic stresses, including DS, lead to an accumulation of misfolded or unfolded proteins, which induces an imbalance in ER homeostasis, a phenomenon known as ER stress (Manghwar & Li, 2022). A cytoprotective response called the UPR (unfolded protein response) is activated to overcome ER stress (Brandizzi *et al*., 2014). The KDEL receptor recovery pathway plays an essential role in this process, by retaining ER-resident proteins (Wires *et al*., 2021). This molecular pathway was found to be upregulated in SO but not in PO and probably helps to counteract the effects of DS.

Finally, we identified a gene encoding a NAD-dependent malate dehydrogenase that plays a key role in the short-term adjustment of stromal NADP(H) redox state in response to changing environments. The maintenance of redox homeostasis is a key molecular mechanism that enables cells to maintain their metabolism under DS (Hebbelmann *et al*., 2012; Kandoi *et al*., 2018).

### Potential signatures of local adaptation in SO and PO

In oak, as in rice (Garg *et al*., 2015), sensitivity to drought and other abiotic stresses seems to be associated with differences in the methylation landscape, even in the absence of stress, suggesting a long term epigenetic adaptation to different ecophysiological situations. Consistently, significant differences in the distribution of DNA methylation were observed between SO and PO, with most of the DMRs identified in CG and CHG contexts (Fig.5). Several studies have suggested that differences in CG and CHG methylation contexts reflect epigenetic variation associated with environmental conditions (Rico *et al*., 2014; Dubin *et al*., 2015; Platt *et al*., 2015; Gugger *et al*., 2016; Browne *et al*., 2021) as in Arabidopsis (Rico *et al*., 2014; Dubin *et al*., 2015; Platt *et al*., 2015; Gugger *et al*., 2016; Browne *et al*., 2021).

Most DMRs with a genomic location in gene bodies, promoters and/or TEs, were hypermethylated in PO relative to SO, in both conditions. Other studies on wheat and rice have reported that hypermethylation is associated with drought-sensitivity (Gayacharan & Joel, 2013; Garg *et al*., 2015; Kaur *et al*., 2018).

The 36315 DMRs (Fig. 4), between PO and SO which were conserved in C and DS conditions and are not related to the response to drought stress were mostly in the CG and CHG context. Genes bearing these stable epigenetic marks were enriched in genetic polymorphisms with large allele frequency differences between populations of the two species (high fixation index FST). Highly differentiated allele frequencies between populations of closely related species include genomic regions contributing to the adaptation of the species to their respective ecological niches, loci involved in reproductive isolation, as well as loci where allele frequencies have diverged due to genetic drift (Le Provost *et al*., 2021). The latter is unlikely because we focus on extremes of the genome-wide FST distribution, and the other two categories are not mutually exclusive, as local adaptation can drive reproductive isolation. In any case, finding that genes bearing these stable epigenetic differences between species often display genetic differentiation between the two species, related to adaptation or reproductive isolation, points to a link between sequence evolution and local patterns of methylations.

The common DMRs are enriched in GO terms related to ‘protein phosphorylation’, ‘signal transduction’, ‘carbohydrate metabolic process’ and ‘cell macromolecule catabolic process’. Some of the genes encode important cytoskeleton proteins, including kinesins and beta-glucosidases. Kinesins are motor proteins involved in cell movement, division, and transport, and in the maintenance of cellular shape (Ali & Yang, 2020). Beta-glucosidases have very diverse roles in plants: cell wall lignification and degradation, activation of several phytohormones and generation of signal molecules (Morant *et al*., 2008). A role for this enzyme in DS responses has also been suggested in Arabidopsis (Han *et al*., 2012) and mulberry (Ackah *et al*., 2022).

In conclusion, some of these DMRs could be used as epigenetic markers with the potential to improve our understanding of population structure and phenotypic variability, as reported by Sow et al., (2018) and may contribute to the development of markers that could be used in the context of -assisted migration (Aitken & Whitlock, 2013) considering the interaction between genetics, epigenetics and environment(Sow *et al*., 2021).

### The linked response between smRNA, DNA methylation and transcriptomic modifications

A comparative analysis of the three data sets generated in this study identified 26 genomic regions in which smRNA clusters were associated with hypermethylation of the corresponding DMR, leading to downregulation of the corresponding gene (Table 1). In this context, these genomic regions may reflect RNA silencing mechanisms in which smRNA clusters guide DNA methylation via transcriptional gene silencing. An alternative interpretation would be that the absence of a smRNA cluster does not lead to DNA hypermethylation, and that overexpression of the corresponding gene may therefore be observed.

Seven of these genomic regions are of particular interest: (i) two encoding basic chitinases proteins, downregulated in SO in C conditions (ii) two encoding proteins that are highly similar to lipoxygenase proteins (LOX) overexpressed in SO in C conditions and (iii) three encoding cytochrome P450 proteins upregulated in SO in DS conditions (Table 1). LOXs have been shown to be involved in the response to DS in olive trees (Sofo *et al*., 2004), pepper (Lim *et al*., 2015)) and *Brassica rapa* (Rai *et al*., 2021). In these studies, an overexpression of LOX genes was generally observed in SO suggesting that its DS tolerance may be related to higher basal levels of these drought-tolerance gene expression.

Finally, P450s constitute the largest family of plant enzymes involved in plant metabolism, including hormone biosynthesis, the synthesis of primary and secondary metabolites, and catabolism. These proteins have recently been implicated in abiotic stress responses and, particularly, in DS response (Pandian *et al*., 2020). The overexpression of P450 genes was reported in several plant species, including maize (Li & Wei, 2020), poplar (Cheng *et al*., 2021) and Arabidopsis (Kushiro *et al*., 2004). Interestingly, P450 is involved in the synthesis of cell wall components. More widely, P450 enzymes are known to be involved in various biosynthetic and detoxification pathways (Pandian *et al*., 2020). In SO, the transcriptomic analysis highlighted an upregulation of several genes associated with these biological processes (NAD-dependent malate dehydrogenases (i.e. detoxification pathways), cytoskeleton components, cellulose synthases and beta-glucosidases (i.e cell-wall remodeling and modification). The overexpression of P450 in SO may, therefore, account for the better DS tolerance of this species.

### General conclusions

We report here the first integrative study to shed light on the ecological preferences of two sympatric oak species (PO and SO) differing in their adaptation to soil water deficit. This experiment was focussed on a relatively short drought stress, starting from just above the level of soil water content at which canopy transpiration is reduced, to just below. This is a critical range in which a different response strategy between the two species might be observed. In accordance with physiological data, the transcriptome reprogramming was stronger in PO than SO suggesting that each species responds in a specific way to DS and that PO is mobilizing more resources than SO to adapt to DS. Similarly, methylome analysis shows that DS induces a species-specific DNA methylation response, mostly in the CHH context, suggesting that this context is the more dynamic under environmental changes. The other two contexts (CG and CHG) displayed differences between PO and SO with a majority of them maintained in both conditions These DMRs might be useful markers to discriminate PO from SO. These DMRs were enriched in highly differentiated SNPs, suggesting that some of these markers may be associated both with the ecological differences or intrinsic barriers to reproduction between the two species. Finally, the overlaps observed between the three -omics datasets strongly suggest that there is a species-specific response to DS. Genomic regions relating to cell-wall processes were identified in SO, which contrast with the wider range of physiological processes mobilized in PO. The cell wall processes may sustain the importance of these biological processes in the DS response, potentially underlying the better tolerance to water deprivation observed in SO. Future studies will investigate the potential role of the genes located in these genomic regions in the adaptation to DS through population genetic analyses (i.e. outlier SNPs loci).

## Author contributions

The work was performed in the frame of the METDRY project. BR analyzed the datasets with PG and wrote the manuscript with GLP and PG. GLP and PG designed and coordinated the research. OB and TG produced the plant material and were involved in the drought stress experiment. BB performed the FST enrichment analysis. All the authors approved the final version of the manuscript. BR and GLP contributed equally to this work. PG coordinated the Metdry Project of the Cluster of Excellence COTE.

## Acknowledgments

The experimental design was supported by the ANR H2oak project (2014 14-CE02-0013-02, “Diversity for adaptive traits related to water use in two temperate European white oaks”). BR was funded both by the Cluster of Excellence COTE (ANR-10-LABX-45) and the EPIDEP project (Plan National du Dépérissement de la Vigne). Sequencing costs were supported by the METDRY project (Cluster of Excellence COTE-ANR-10-LABX-45).

## Data availability

All the sequences used in this study have been deposited in the National Center for Biotechnology Information (NCBI) Sequence Read Archive https://www.ncbi.nlm.nih.gov/sra) under accession number PRJNA810970.

## References

Ackah M, Guo L, Li S, Jin X, Asakiya C, Aboagye ET, Yuan F, Wu M, Essoh LG, Adjibolosoo D, et al. 2022. DNA Methylation Changes and Its Associated Genes in Mulberry (Morus alba L.) Yu-711 Response to Drought Stress Using MethylRAD Sequencing. Plants 11.

Aitken SN, Whitlock MC. 2013. Assisted gene flow to facilitate local adaptation to climate change. Annual review of ecology, evolution, and systematics 44: 367–388.

Alexa A. 2006. Gene set enrichment analysis with topGO.

Ali I, Yang W-C. 2020. The functions of kinesin and kinesin-related proteins in eukaryotes. Cell adhesion & migration 14: 139–152.

Allimuthu E, Dalal M, Kumar KG, Sellathdurai D, Kumar RR, Sathee L, Chinnusamy V. 2020. Characterization of Atypical Protein Tyrosine Kinase (PTK) Genes and Their Role in Abiotic Stress Response in Rice. Plants 9.

Arcuri MLC, Fialho LC, Vasconcellos Nunes-Laitz A, Fuchs-Ferraz MCP, Wolf IR, Valente GT, Marino CL, Maia IG. 2020. Genome-wide identification of multifunctional laccase gene family in Eucalyptus grandis: potential targets for lignin engineering and stress tolerance. Trees 34: 745–758.

Arneth, Harrison, Zaehle, Tsigaridis. Terrestrial biogeochemical feedbacks in the climate system. Nature.

Axtell MJ. 2013. ShortStack: comprehensive annotation and quantification of small RNA genes. RNA 19: 740–751.

Behringer D, Zimmermann H, Ziegenhagen B, Liepelt S. 2015. Differential Gene Expression Reveals Candidate Genes for Drought Stress Response in Abies alba (Pinaceae). PloS one 10: e0124564.

Bogeat-Triboulot MB, Buré C, Gerardin T, Chuste PA, Le Thiec D, Hummel I, Durand M, Wildhagen H, Douthe C, Molins A, et al. 2019. Additive effects of high growth rate and low transpiration rate drive differences in whole plant transpiration efficiency among black poplar genotypes. Environmental and experimental botany 166: 103784.

Brandizzi F, Frigerio L, Howell SH, Schäfer P. 2014. Endoplasmic reticulum— shape and function in stress translation. Frontiers in plant science 5.

Browne L, MacDonald B, Fitz-Gibbon S, Wright JW, Sork VL. 2021. Genome-Wide Variation in DNA Methylation Predicts Variation in Leaf Traits in an Ecosystem- Foundational Oak Species. *Forests*, Trees and Livelihoods 12: 569.

Caboche S, Audebert C, Lemoine Y, Hot D. 2014. Comparison of mapping algorithms used in high-throughput sequencing: application to Ion Torrent data. BMC genomics 15: 264.

Ceccherini G, Duveiller G, Grassi G, Lemoine G, Avitabile V, Pilli R, Cescatti A. 2020. Abrupt increase in harvested forest area over Europe after 2015. Nature 583: 72–77.

Cheng H, Shao Z, Lu C, Duan D. 2021. Genome-wide identification of chitinase genes in Thalassiosira pseudonana and analysis of their expression under abiotic stresses. BMC plant biology 21: 87.

Daviaud C, Renault V, Mauger F, Deleuze J-F, Tost J. 2018. Whole-Genome Bisulfite Sequencing Using the Ovation® Ultralow Methyl-Seq Protocol. Methods in molecular biology 1708: 83–104.

Dubin MJ, Zhang P, Meng D, Remigereau M-S, Osborne EJ, Paolo Casale F, Drewe P, Kahles A, Jean G, Vilhjálmsson B, et al. 2015. DNA methylation in Arabidopsis has a genetic basis and shows evidence of local adaptation. eLife 4: e05255.

Du M, Ding G, Cai Q. 2018. The Transcriptomic Responses of Pinus massoniana to Drought Stress. Forests 9: 326.

Eaton, Caudullo, Oliveira. 2016. Quercus robur and Quercus petraea in Europe: distribution, habitat, usage and threats. European atlas of forest.

Epron D, Dreyer E. 1990. Stomatal and non stomatal limitation of photosynthesis by leaf water deficits in three oak species: a comparison of gas exchange and chlorophyll a fluorescence data. Annales des Sciences Forestières 47: 435–450.

Erdmann RM, Picard CL. 2020. RNA-directed DNA Methylation. PLoS genetics 16: e1009034.

Estravis-Barcala M, Mattera MG, Soliani C, Bellora N, Opgenoorth L, Heer K, Arana MV. 2020. Molecular bases of responses to abiotic stress in trees. Journal of experimental botany 71: 3765–3779.

Ezquer I, Salameh I, Colombo L, Kalaitzis P. 2020. Plant Cell Walls Tackling Climate Change: Biotechnological Strategies to Improve Crop Adaptations and Photosynthesis in Response to Global Warming. Plants 9: 212.

Feng H, Wu H. 2019. Differential methylation analysis for bisulfite sequencing using DSS. *Quantitative biology (Beijing*, China) 7: 327–334.

Fox H, Doron-Faigenboim A, Kelly G, Bourstein R, Attia Z, Zhou J, Moshe Y, Moshelion M, David-Schwartz R. 2018. Transcriptome analysis of Pinus halepensis under drought stress and during recovery. Tree physiology 38: 423–441.

Gao X, Ren F, Lu Y-T. 2006. The Arabidopsis mutant stg1 identifies a function for TBP-associated factor 10 in plant osmotic stress adaptation. Plant & cell physiology 47: 1285–1294.

Garg R, Narayana Chevala V, Shankar R, Jain M. 2015. Divergent DNA methylation patterns associated with gene expression in rice cultivars with contrasting drought and salinity stress response. Scientific reports 5: 14922.

Gayacharan, Joel AJ. 2013. Epigenetic responses to drought stress in rice (Oryza sativa L.). Physiology and molecular biology of plants: an international journal of functional plant biology 19: 379–387.

Granier A, Loustau D, Bréda N. 2000. A generic model of forest canopy conductance dependent on climate, soil water availability and leaf area index. Annals of forest science 57: 755–765.

Guerrero-Sánchez VM, Castillejo MÁ, López-Hidalgo C, Alconada AMM, Jorrín-Novo JV, Rey M-D. 2021. Changes in the transcript and protein profiles of Quercus ilex seedlings in response to drought stress. Journal of proteomics 243: 104263.

Gugger PF, Fitz-Gibbon S, PellEgrini M, Sork VL. 2016. Species-wide patterns of DNA methylation variation in Quercus lobata and their association with climate gradients. Molecular ecology 25: 1665–1680.

Gugger PF, Peñaloza-Ramírez JM, Wright JW, Sork VL. 2017. Whole-transcriptome response to water stress in a California endemic oak, Quercus lobata. Tree physiology 37: 632–644.

Han Y-J, Cho K-C, Hwang O-J, Choi Y-S, Shin A-Y, Hwang I, Kim J-I. 2012. Overexpression of an Arabidopsis β-glucosidase gene enhances drought resistance with dwarf phenotype in creeping bentgrass. Plant cell reports 31: 1677–1686.

Haneca K, Katarina Čufar, Beeckman H. 2009. Oaks, tree-rings and wooden cultural heritage: a review of the main characteristics and applications of oak dendrochronology in Europe. Journal of archaeological science 36: 1–11.

Hanewinkel M, Cullmann DA, Schelhaas M-J, Nabuurs G-J, Zimmermann NE. 2012. Climate change may cause severe loss in the economic value of European forest land. Nature climate change 3: 203–207.

He L, Huang H, Bradai M, Zha, You Y, Ma J, Zao L, Lozano Duran R, Zhu JK. 2022. DNA methylation-free *Arabidopsis* reveals crucial roles of DNA methylation in regulating gene expression and development. Nature Communications 13:1335–1356.

Hebbelmann I, Selinski J, Wehmeyer C, Goss T, Voss I, Mulo P, Kangasjärvi S, Aro E-M, Oelze M-L, Dietz K-J, et al. 2012. Multiple strategies to prevent oxidative stress in Arabidopsis plants lacking the malate valve enzyme NADP-malate dehydrogenase. Journal of experimental botany 63: 1445–1459.

Kandoi D, Mohanty S, Tripathy BC. 2018. Overexpression of plastidic maize NADP-malate dehydrogenase (ZmNADP-MDH) in Arabidopsis thaliana confers tolerance to salt stress. Protoplasma 255: 547–563.

Kaur A, Grewal A, Sharma P. 2018. Comparative analysis of DNA methylation changes in two contrasting wheat genotypes under water deficit. Biologia plantarum 62: 471–478.

Kesten C, Menna A, Sánchez-Rodríguez C. 2017. Regulation of cellulose synthesis in response to stress. Current opinion in plant biology 40: 106–113.

Krueger F, Andrews SR. 2011. Bismark: a flexible aligner and methylation caller for Bisulfite-Seq applications. Bioinformatics 27: 1571–1572.

Kumar S, Mohapatra T. 2021. Dynamics of DNA Methylation and Its Functions in Plant Growth and Development. Frontiers in plant science 12: 596236.

Kushiro T, Okamoto M, Nakabayashi K, Yamagishi K, Kitamura S, Asami T, Hirai N, Koshiba T, Kamiya Y, Nambara E. 2004. The Arabidopsis cytochrome P450 CYP707A encodes ABA 8’-hydroxylases: key enzymes in ABA catabolism. The EMBO journal 23: 1647–1656.

Lafon-Placette C, Le Gac A-L, Chauveau D, Segura V, Delaunay A, Lesage-Descauses M-C, Hummel I, Cohen D, Jesson B, Le Thiec D, et al. 2018. Changes in the epigenome and transcriptome of the poplar shoot apical meristem in response to water availability affect preferentially hormone pathways. Journal of experimental botany 69: 537–551.

Lamka, Harder, Sundaram. Epigenetics in ecology, evolution, and conservation. Frontiers in ecology and evolution.

Le Gall H, Philippe F, Domon J-M, Gillet F, Pelloux J, Rayon C. 2015. Cell Wall Metabolism in Response to Abiotic Stress. Plants 4: 112–166.

Le Provost G, Brachi B, Lesur I, Lalanne C, Labadie K, Aury J-M, Da Silva C, Leroy T, Plomion C. 2021. Water stress-associated isolation barriers between two sympatric oak species. bioRxiv: 2021.09.16.460585.

Le Provost G, Gerardin T, Plomion C, Brendel O. 2022. Molecular plasticity to soil water deficit differs between sessile oak (Quercus Petraea (Matt.) Liebl.) high- and low-water use efficiency genotypes. Tree physiology.

Liang D, Zhang Z, Wu H, Huang C, Shuai P, Ye C-Y, Tang S, Wang Y, Yang L, Wang J, et al. 2014. Single-base-resolution methylomes of populus trichocarpa reveal the association between DNA methylation and drought stress. BMC Genetics 15.

Liao Y, Smyth GK, Shi W. 2014. featureCounts: an efficient general purpose program for assigning sequence reads to genomic features. Bioinformatics 30: 923– 930.

Lim CW, Han S-W, Hwang IS, Kim DS, Hwang BK, Lee SC. 2015. The Pepper Lipoxygenase CaLOX1 Plays a Role in Osmotic, Drought and High Salinity Stress Response. Plant & cell physiology 56: 930–942.

Li Y, Wei K. 2020. Comparative functional genomics analysis of cytochrome P450 gene superfamily in wheat and maize. BMC plant biology 20: 93.

Love, Anders, Huber. 2014. Differential analysis of count data–the DESeq2 package. Genome biology.

Lu X, Wang X, Chen X, Shu N, Wang J, Wang D, Wang S, Fan W, Guo L, Guo X, et al. 2017. Single-base resolution methylomes of upland cotton (Gossypium hirsutum L.) reveal epigenome modifications in response to drought stress. BMC genomics 18: 297.

Madritsch S, Wischnitzki E, Kotrade P, Ashoub A, Burg A, Fluch S, Brüggemann W, Sehr EM. 2019. Elucidating Drought Stress Tolerance in European Oaks Through Cross-Species Transcriptomics. G3 9: 3181–3199.

Ma H, Liu M. 2019. The microtubule cytoskeleton acts as a sensor for stress response signaling in plants. Molecular biology reports 46: 5603–5608.

Manghwar H, Li J. 2022. Endoplasmic Reticulum Stress and Unfolded Protein Response Signaling in Plants. International journal of molecular sciences 23.

Morant AV, Jørgensen K, Jørgensen C, Paquette SM, Sánchez-Pérez R, Møller BL, Bak S. 2008. beta-Glucosidases as detonators of plant chemical defense. Phytochemistry 69: 1795–1813.

Müller M, Seifert S, Lübbe T, Leuschner C, Finkeldey R. 2017. De novo transcriptome assembly and analysis of differential gene expression in response to drought in European beech. PloS one 12: e0184167.

Mun B-G, Lee S-U, Park E-J, Kim H-H, Hussain A, Imran QM, Lee I-J, Yun B-W. 2017. Analysis of transcription factors among differentially expressed genes induced by drought stress in Populus davidiana. 3 Biotech 7: 209.

Nadarajah KK. 2020. ROS Homeostasis in Abiotic Stress Tolerance in Plants. International journal of molecular sciences 21.

Niederhuth CE, Bewick AJ, Ji L, Alabady MS, Kim KD, Li Q, Rohr NA, Rambani A, Burke JM, Udall JA, et al. 2016. Widespread natural variation of DNA methylation within angiosperms. Genome biology 17: 194.

Pandian BA, Sathishraj R, Djanaguiraman M, Prasad PVV, Jugulam M. 2020. Role of Cytochrome P450 Enzymes in Plant Stress Response. *Antioxidants (Basel*, Switzerland) 9.

Parvathi MS, Nataraja KN, Nanja Reddy YA, Naika MBN, Channabyre Gowda MV. 2019. Transcriptome analysis of finger millet (Eleusine coracana (L.) Gaertn.) reveals unique drought responsive genes. Journal of genetics 98.

Platt A, Gugger PF, Pellegrini M, Sork VL. 2015. Genome-wide signature of local adaptation linked to variable CpG methylation in oak populations. Molecular ecology 24: 3823–3830.

Plomion C, Aury J-M, Amselem J, Alaeitabar T, Barbe V, Belser C, Bergès H, Bodénès C, Boudet N, Boury C, et al. 2016. Decoding the oak genome: public release of sequence data, assembly, annotation and publication strategies. Molecular ecology resources 16: 254–265.

Ponton S, Dupouey J-L, Bréda N, Dreyer E. 2002. Comparison of water-use efficiency of seedlings from two sympatric oak species: genotype x environment interactions. Tree physiology 22: 413–422.

Ponton S, Dupouey J-L, Breda N, Feuillat F, Bodenes C, Dreyer E. 2001. Carbon isotope discrimination and wood anatomy variations in mixed stands of Quercus robur and Quercus petraea. Plant, cell & environment 24: 861–868.

Puranik S, Sahu PP, Srivastava PS, Prasad M. 2012. NAC proteins: regulation and role in stress tolerance. Trends in plant science 17: 369–381.

Quinlan AR, Hall IM. 2010. BEDTools: a flexible suite of utilities for comparing genomic features. Bioinformatics 26: 841–842.

Rai AN, Mandliya T, Kulkarni P, Rao M, Suprasanna P. 2021. Evolution and Transcriptional Modulation of Lipoxygenase Genes Under Heat, Drought, and Combined Stress in Brassica rapa. Plant molecular biology reporter / ISPMB 39: 60– 71.

Richards CL, Bossdorf O, Pigliucci M. 2010. What Role Does Heritable Epigenetic Variation Play in Phenotypic Evolution? Bioscience 60: 232–237.

Richardson AD, Hufkens K, Milliman T, Aubrecht DM, Furze ME, Seyednasrollah B, Krassovski MB, Latimer JM, Nettles WR, Heiderman RR, et al. 2018. Ecosystem warming extends vegetation activity but heightens vulnerability to cold temperatures. Nature 560: 368–371.

Rico L, Ogaya R, Barbeta A, Peñuelas J. 2014. Changes in DNA methylation fingerprint of Quercus ilex trees in response to experimental field drought simulating projected climate change. Plant biology 16: 419–427.

Rodríguez, Canales. 2005. Molecular aspects of abiotic stress in plants. Biotecnologia aplicada: revista de la Sociedad Iberolatinoamericana para Investigaciones sobre Interferon y Biotecnologia en Salud.

Rubio B, Stammitti L, Cookson SJ, Teyssier E, Gallusci P. 2022. Small RNA populations reflect the complex dialogue established between heterograft partners in grapevine. Horticulture research.

Schmull M, Thomas FM. 2000. Morphological and physiological reactions of young deciduous trees (Quercus robur L., Q. petraea [Matt.] Liebl., Fagus sylvatica L.) to waterlogging. Plant and soil 225: 227–242.

Sharma, Jha, Dubey. 2012. Reactive oxygen species, oxidative damage, and antioxidative defense mechanism in plants under stressful conditions. Journal of atomic and molecular physics.

Singh D, Laxmi A. 2015. Transcriptional regulation of drought response: a tortuous network of transcriptional factors. Frontiers in plant science 6: 895.

Soda N, Singla-Pareek SL, Pareek A. 2016. Abiotic stress response in plants: Role of cytoskeleton. In: Abiotic Stress Response in Plants. Weinheim, Germany: Wiley-VCH Verlag GmbH & Co. KGaA, 107–134.

Sofo A, Dichio B, Xiloyannis C, Masia A. 2004. Lipoxygenase activity and proline accumulation in leaves and roots of olive trees in response to drought stress. Physiologia plantarum 121: 58–65.

Sork VL. 2017. Genomic Studies of Local Adaptation in Natural Plant Populations. The Journal of heredity 109: 3–15.

Sow MD, Le Gac A-L, Fichot R, Lanciano S, Delaunay A, Le Jan I, Lesage-Descauses M-C, Citerne S, Caius J, Brunaud V, et al. 2021. RNAi suppression of DNA methylation affects the drought stress response and genome integrity in transgenic poplar. The New phytologist 232: 80–97.

Spinoni J, Vogt JV, Naumann G, Barbosa P, Dosio A. 2018. Will drought events become more frequent and severe in Europe? International Journal of Climatology 38: 1718–1736.

Tang S, Dong Y, Liang D, Zhang Z, Ye C-Y, Shuai P, Han X, Zhao Y, Yin W, Xia X. 2015. Analysis of the Drought Stress-Responsive Transcriptome of Black Cottonwood (Populus trichocarpa) Using Deep RNA Sequencing. Plant molecular biology reporter / ISPMB 33: 424–438.

Torre S, Tattini M, Brunetti C, Fineschi S, Fini A, Ferrini F, Sebastiani F. 2014. RNA-seq analysis of Quercus pubescens Leaves: de novo transcriptome assembly, annotation and functional markers development. PloS one 9: e112487.

Tweneboah S, Oh S-K. 2017. Biological roles of NAC transcription factors in the regulation of biotic and abiotic stress responses in solanaceous crops. Journal of Plant Biotechnology 44: 1–11.

Villar E, Klopp C, Noirot C, Novaes E, Kirst M, Plomion C, Gion J-M. 2011. RNA-Seq reveals genotype-specific molecular responses to water deficit in eucalyptus. BMC genomics 12: 538.

Wang T, McFarlane HE, Persson S. 2016. The impact of abiotic factors on cellulose synthesis. Journal of experimental botany 67: 543–552.

Wires ES, Trychta KA, Kennedy LM, Harvey BK. 2021. The Function of KDEL Receptors as UPR Genes in Disease. International journal of molecular sciences 22.

Yamamoto K, Hamada H, Shinkai H, Kohno Y, Koseki H, Aoe T. 2003. The KDEL receptor modulates the endoplasmic reticulum stress response through mitogen-activated protein kinase signaling cascades. The Journal of biological chemistry 278: 34525–34532.

Zhang H, Lang Z, Zhu J-K. 2018. Dynamics and function of DNA methylation in plants. Nature Reviews Molecular Cell Biology 19: 489–506.

Zhang Y, Zhao L, Xiao H, Chew J, Xiang J, Qian K, Fan X. 2020. Knockdown of a Novel Gene OsTBP2.2 Increases Sensitivity to Drought Stress in Rice. Genes 11.

Zhao P, Ma B, Cai C, Xu J. 2022. Transcriptome and methylome changes in two contrasting mungbean genotypes in response to drought stress. BMC genomics 23: 80.

