## Supplementary information (Tables and Figures) for "Species-specific epigenetic responses to drought stress of two sympatric oak species reflect their ecological preferences"

**Methods S1** Additional details for the experimental design.

PO and SO seedlings grown in greenhouses at the National Institute of Agronomic Research (INRA), Champenoux, France (48°45’8’’N, 6°20’28’’E, 259m). The acorns used to generate the seedlings for this experiment were collected in the fall of 2015 from mature PO and SO trees growing in pure stands of these species in France. The PO stand was located in north-eastern France (49°16'43"N 5°28'43"E), at an elevation of 223 m, in an environment classified as moderately humid. The SO stand was located in north-central France (47°32'23"N 1°30'01"E) at an elevation of 93 m, in an environment classified as dry. The seeds were sown in spring 2016, in 1 L pots filled with a mixture of 80% soil (a mixture of 50% topsoil and 50% flax compost) and 20% river sand, which were placed in a shaded common garden. Six seedlings from each provenance (6 plants x 2 provenances x 2 species = 24 plants in total) were transferred to 10 L pots containing a 5:3:2 (by volume) mixture of sand, peat and silty-argillaceous forest soil and fertilized with 15 g Nitrocote per pot. This soil mixture was tested to a field capacity (FC) of 35.0%±1.4% SE volumetric soil water content (SWC; mean of all pots) by time-domain reflectometry (TRIME-TDR; IMKO GmbH, Ettlingen, DE). The experiment took place during six months from April to August 2017 in a greenhouse equipped with a robotic system for the automatic weighing and watering of plants (Bogeat-Triboulot et al. 2019). The air temperature inside the greenhouse followed the environmental variations while never exceeding 25°C due to a cooling system in the facility. The plants were positioned alternatively by species in the greenhouse on day 114 and first submitted to non-limiting growing conditions: natural growth light, well fertilized and irrigated. After an acclimation phase, from day 132 onwards, irrigation was decreased to 70% of the field capacity (REW) to avoid water logging. The relative extractable water was calculated as REW= ((SWC -Wp) / (Fc – Wp))*100 where SWC is the volumetric soil water content, Wp is the wilting point assumed to be at 4% and Fc is the swc at field capacity of each pot/plant . The drought experiment started on day 132, when all the plants were progressively submitted to increasing, stabilised drought levels (Figure 1). Theoretical levels aimed at were 70%- 60%-40%-35%- REW. We selected a subset of three plants per species at random for this study to perform the molecular analysis at C and DS conditions.

**Methods S2** Monitoring of growth and gas exchange.

The diameter (D) of the stem was measured regularly (at two- to three-day intervals) 2 cm above the root collar throughout the entire experiment. Monitoring began in late April (115th day of the year, DoY) and ended in mid-July (205th DoY). The diameter growth rate (DGR) was also calculated from date to date as DGR= (D+1-D) / (t+1 -t), where D is the diameter at a given date, D+1 is the diameter at the following date, t is the DoY at a given date, and t+1 is the DoY at the following date.Gas exchange was monitored with a portable photosynthesis system (LI-COR 6200; LI-COR, Lincoln, NE, USA). CO2 net assimilation rate (An) and stomatal conductance for water vapor (gs) were measured regularly, for each REW level (70, 60, 40, 35% REW) throughout the entire experiment. For each plant, measurements were repeated on the same fully sunlit third-flush leaf grown in the absence of stress. Monitoring began in late May (143th DoY) and ended in mid-July (198th DoY).

**Methods S3** Description of RNA, smRNA and WGBS libraries constrcution and sequencing. RNA, small RNAs and genomic DNA were extracted using 100 mg powder with the mirVana RNA isolation kit (Thermo Fisher Scientific, USA) and the DNeasy plant minikit (Qiagen, Hilden, Germany) according to manufacturer’s instructions. The RNA (total and small-RNA fractions) was quantified on a Nanodrop 1000 spectrophotometer (Thermo Fisher Scientific, USA) and their integrity was assessed (Bioanalyser 2100, Agilent, USA). We selected RNA samples with a minimum RIN of 7 for RNAseq analysis. For library preparation, the mRNA fraction was purified from 5 µg total RNA with Oligo d(T)25 Magnetic Beads from New England Biolabs (Ipswich, MA, USA) according to the manufacturer's instructions. The sequencing was performed with the Ion Torrent Total RNA-seq version2 Kit (Thermo Fisher Scientific, USA, MA), according to the manufacturer’s instructions. Barcoded sequencing adapters were used for the multiplexed sequencing of all 12 sample libraries for small RNAs, and three samples were sequenced simultaneously for gene quantification. Thus, only one flow cell was used for the small-RNA samples, whereas three flow cells were used for transcriptome analysis. High-throughput sequencing was performed with the Ion Proton sequencer available at The Bordeaux Transcriptome Genome Facility (https://pgtb.cgfb.u-bordeaux.fr/). The 12 libraries for WGBS were generated as previously described by Daviaud et al., 2018 with the Ovation Ultralow Methyl-seq kit (Tecan Genomics/ Nugen, San Carlos, CA, USA, http://www.nugen.com/products/ovation-ultralow-methyl-seq-library-systems). The 12 samples to be sequenced were combined, at equimolar concentrations, in a single pool and paired-end sequencing (2 x 150 bp) was performed on four lanes of an Illumina HiSeq4000 flow cell (to yield a minimal theoretical coverage of 30 X for each sample. The Fastq files for each sample were concatenated after sequencing.

**Fig. S1** Monitoring of (a) the mean REW and (b) the mean relative diameter growth rate (DGR) (a) The mean REW is shown for the complete experiment (DoY 115-205) The REW of the plants selected for this experiment is shown for SO in red to the left of the mean and for PO in green to the right of the mean; theoretical REW levels are indicated by vertical grey dashed lines; the two harvests for the genetic analyses are indicated by red dashed lines (b) The DGR is shown for each species and only for the plants selected for this experiment with its standard error of the mean (to indicate the precision of the mean). Significant differences in DGR were tested by t-test and p<0.05 are indicated by *

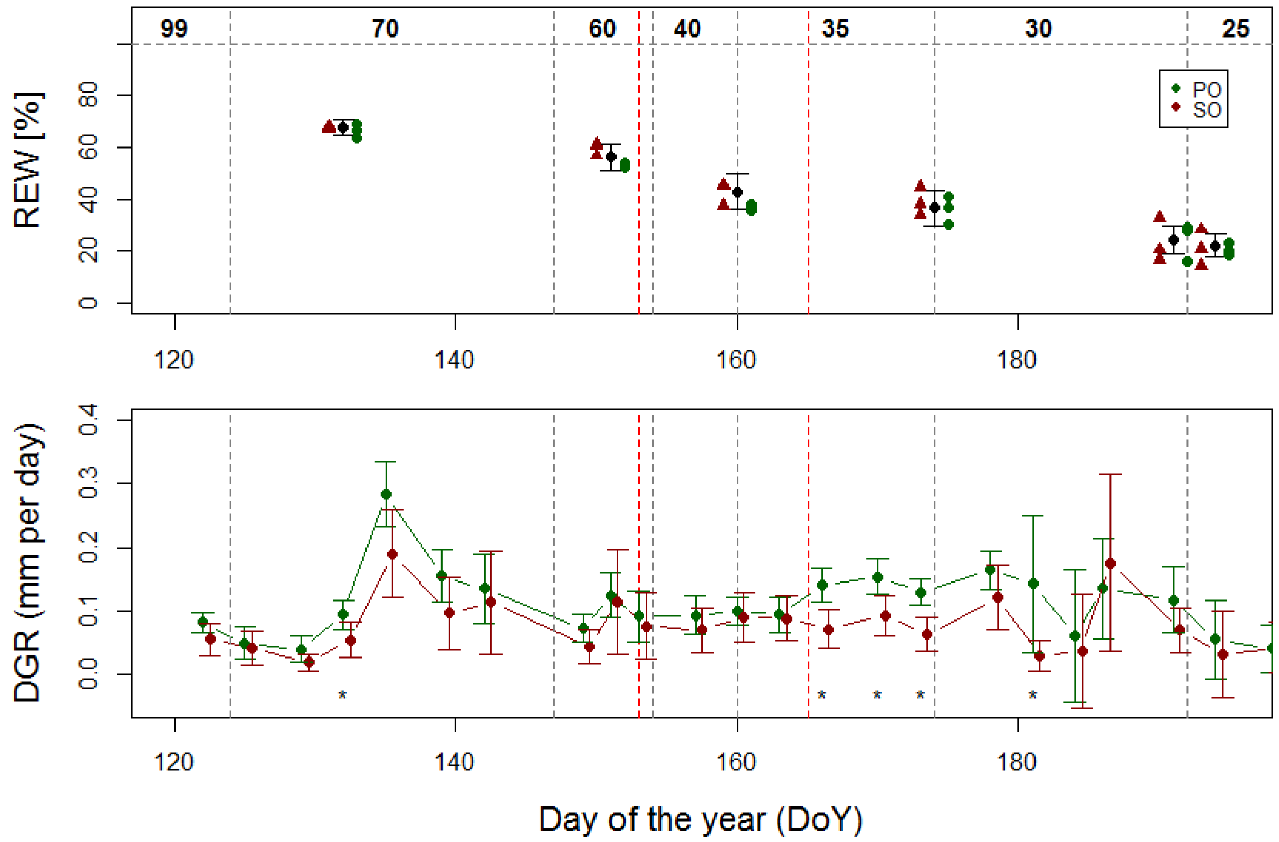

**Fig. S2** Means and standard error of the mean for net CO2 assimilation rate (A), stomatal conductance for water vapour (gs) and intrinsic water use efficiency (WUE) Theoretical REW levels are indicated by vertical grey dashed lines; the two harvests for the geneticanalyses are indicated by red dashed lines. PO and SO are represented by red and green dots respectively. The differences between species at a given date from a t-test are presented as : P values : “***” for P<0.001 ; ”**” for P<0.01 and ”*” for P<0.05.

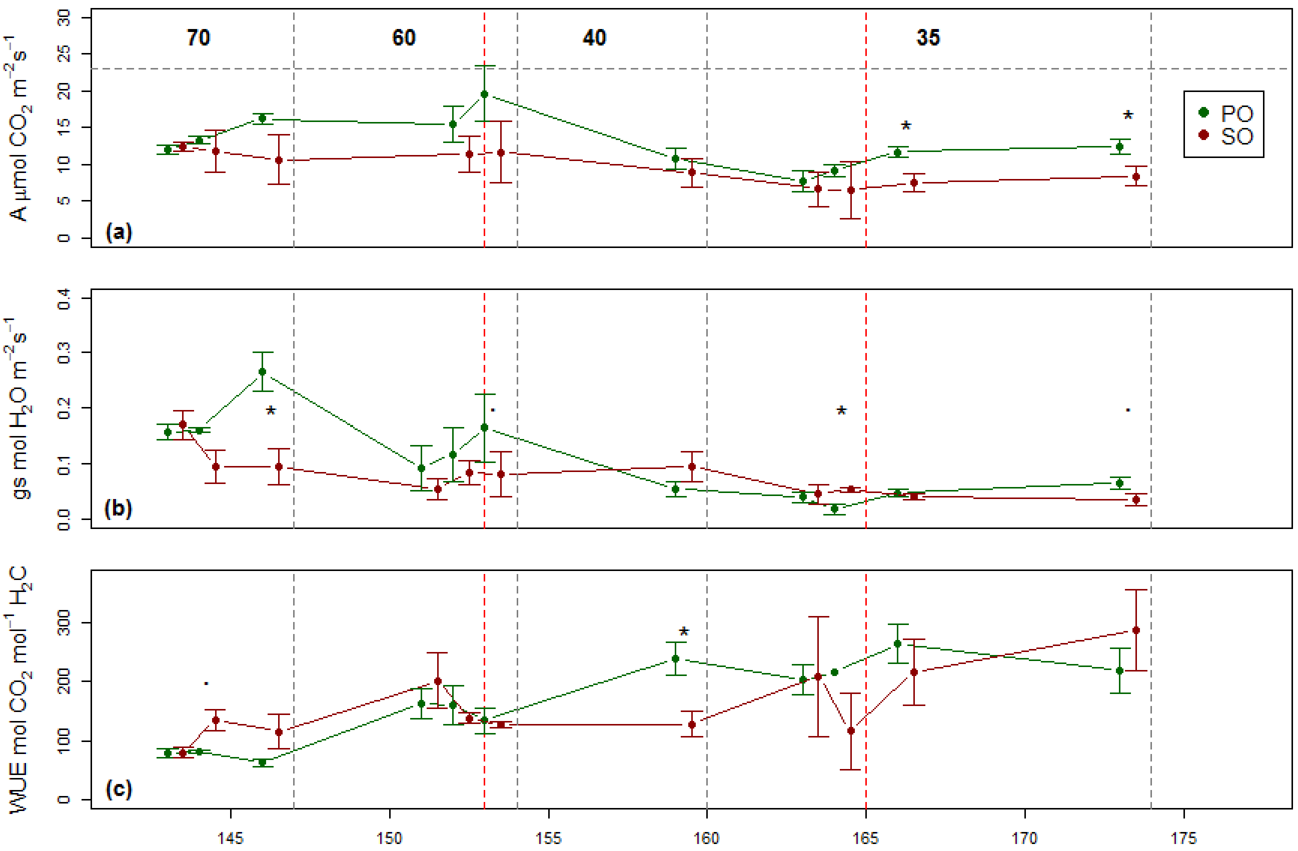

**Fig. S3** Principal component analysis of regularized log-transformed gene expression data. The C and DS samples are presented by the blue and red points, respectively. The SO and PO samples are marked with orange and blue circles, respectively

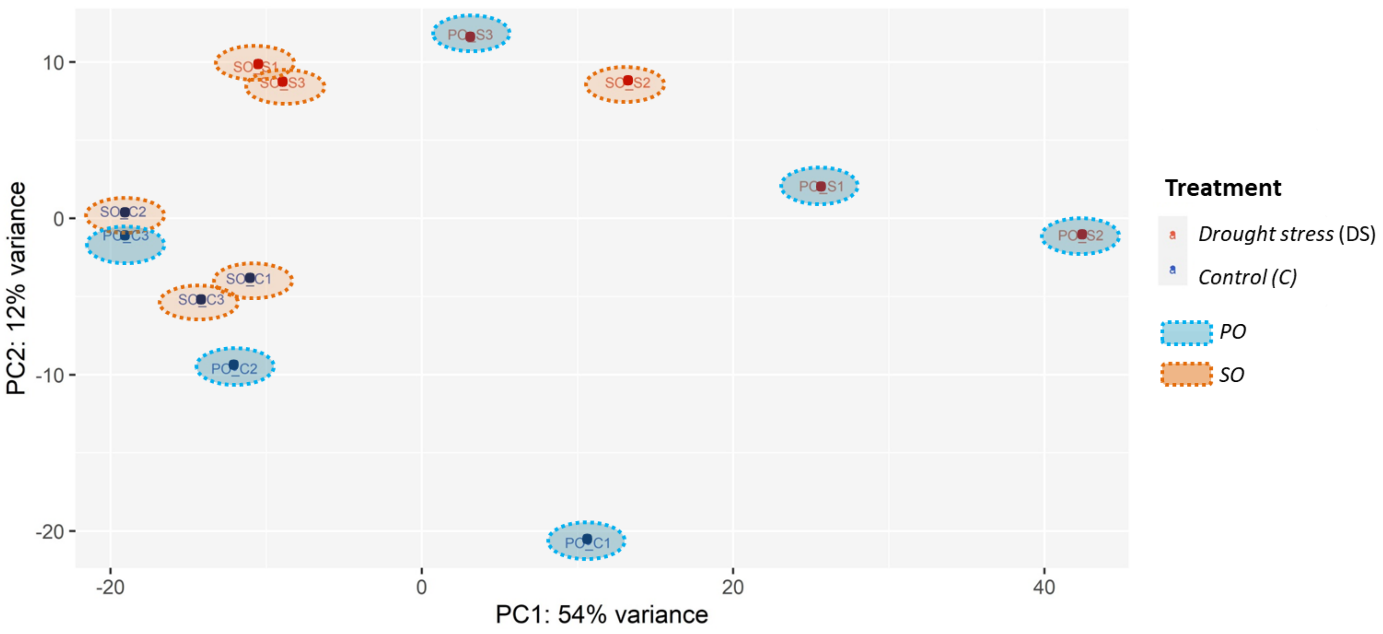


**Fig. S4** DNA methylation levels in CG, CHG and CHH context in PO and SO in C and DS conditions

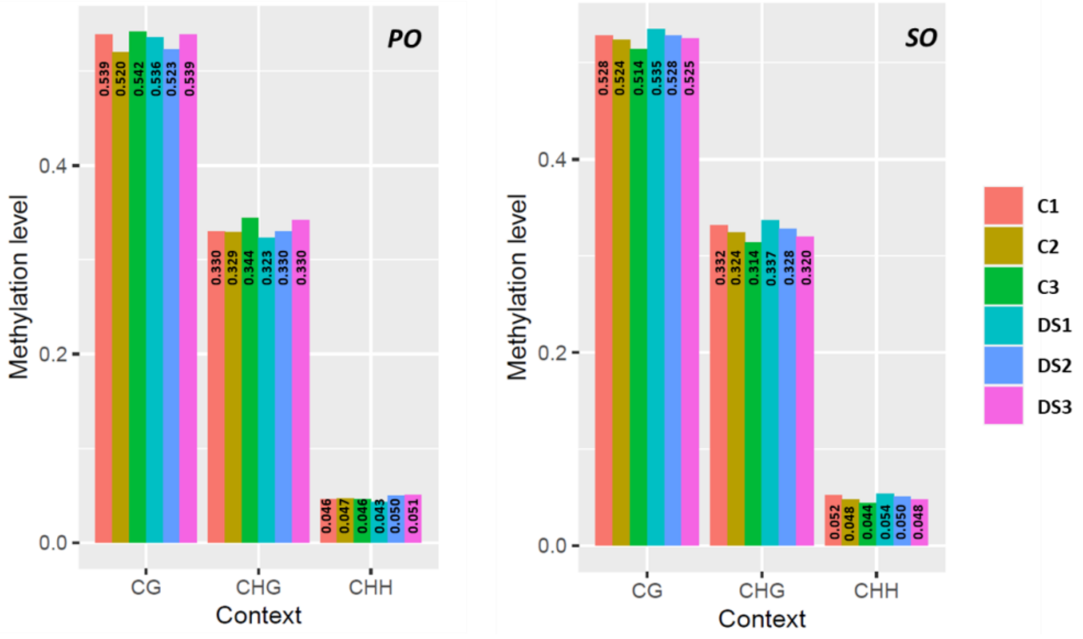

**Fig. S5** The global pattern of SO and PO DNA methylomes in C and DS conditions for CG, CHG and CHH contexts.

The three replicates of each samples are represented. The y-axis indicates the percentage of methylated cytosines according to each methylation level which shown on x-axis

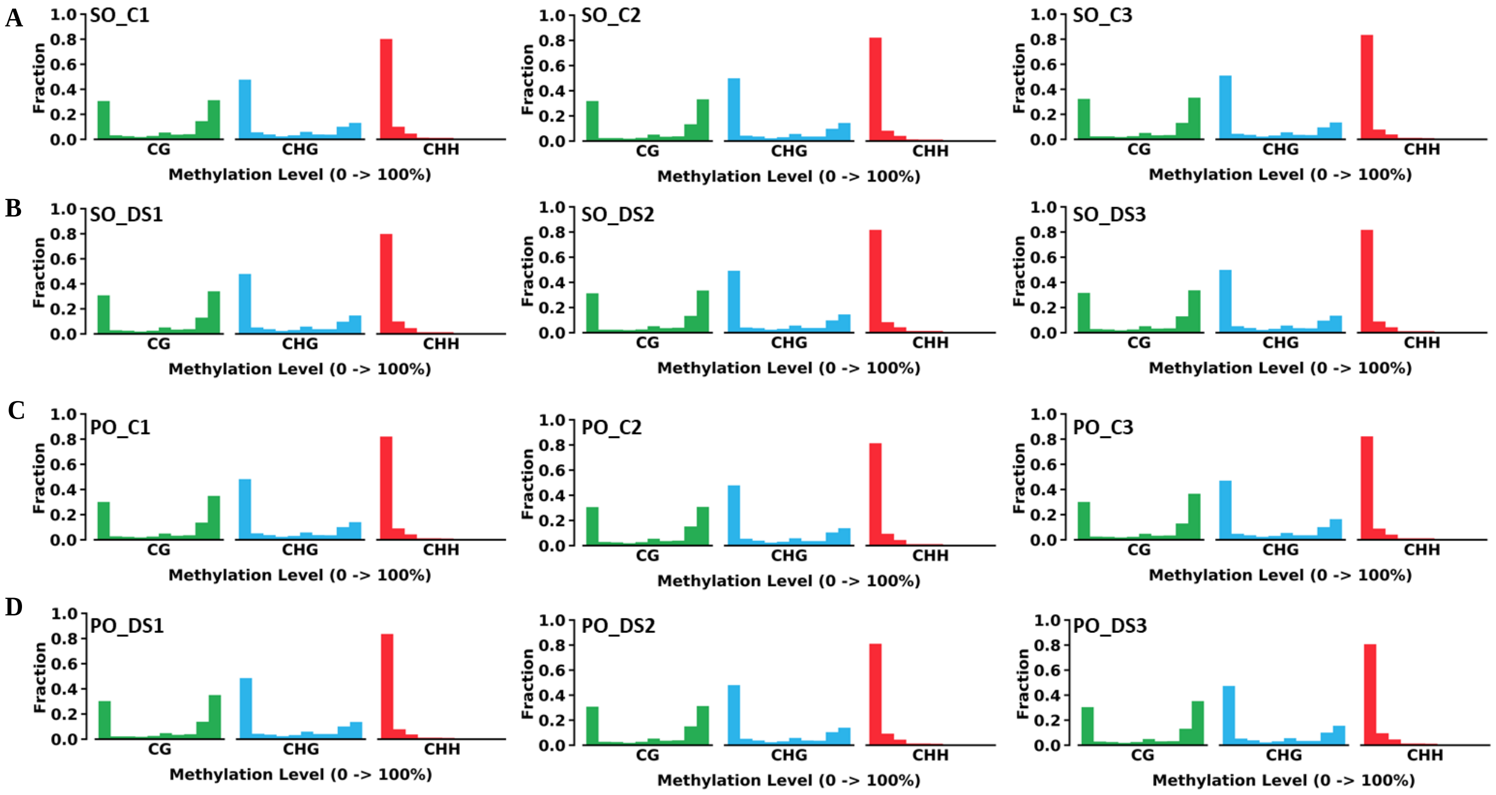

**Fig. S6** DNA methylation pattern in gene bodies and TE regions in PO (A and B) and SO (C and D) samples

The relative fraction of methylated cytosines are represented for control (C) and drought stress (DS) conditions TSS : transcriptional start site ; TTS : transcriptional termination site For annotated genes, we considered seperatly the promoter region 2 kb upstream from the transcriptional start site (TSS), the transcribed region and the region 2 kb downstream from the termination site (TTS). The CG methylation levels were close to 30-35% upstream and downstream from the TSS and TTS of genes. These levels increased within the transcribed region (TR) to reach a maximum of 40%, with no significant difference between species or conditions (A and C). A similar trend was observed in the CHG context, with no difference in the general methylation profile characterized by a mean of 20% methylation downstream and upstream from genes and of almost zero within genes. In this context, the level of methylation within genes was significantly higher (p-value < 0.05) in DS than C conditions, but only in SO (A and C). In the CHH context, methylation levels were very low within genes, increasing to 0.04% in promoters and terminators in both species. Within-gene methylation was higher in C than in DS conditions in PO, whereas the opposite pattern was observed in SO (A and C). Similar results were obtained for comparisons of DNA methylation of TEs between C and DS conditions, except for the CHH context, in which methylation levels increased significantly (p-value < 0.001) in DS conditions for both species (B and D)

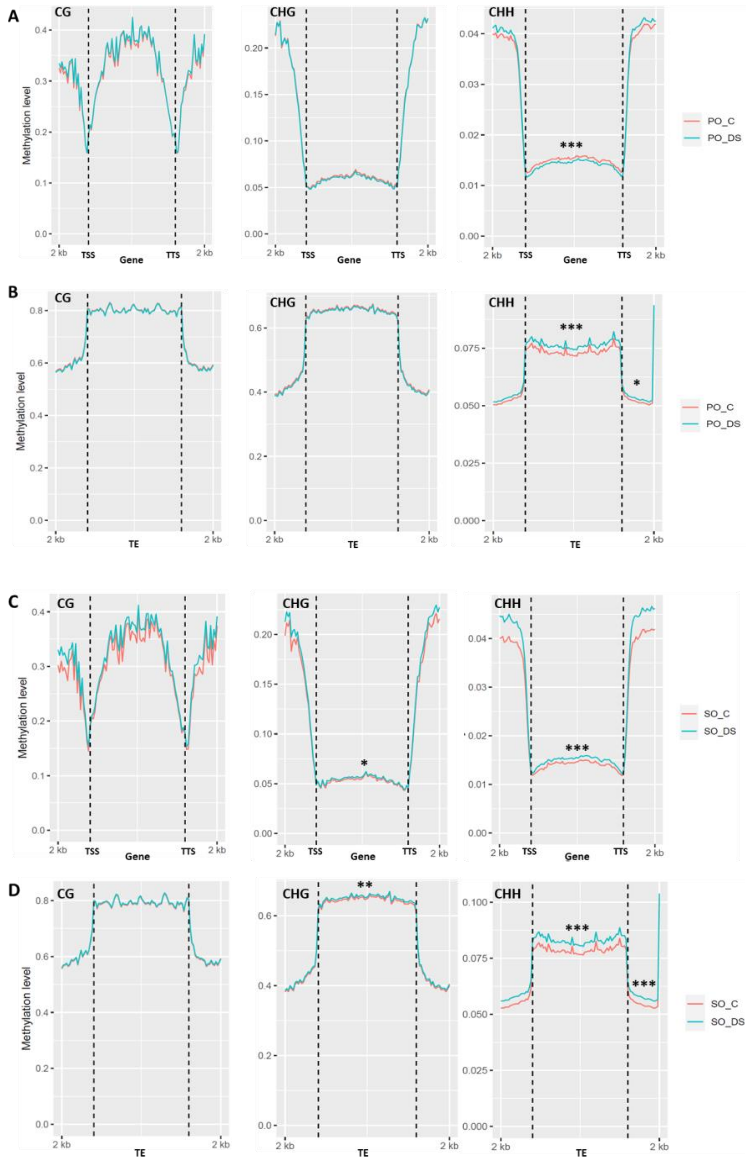

**Fig. S7** Principal component analysis based on the percentage methylation profiles of SO (in red) and PO (in blue) samples in control (C) and drought stress (DS) conditions

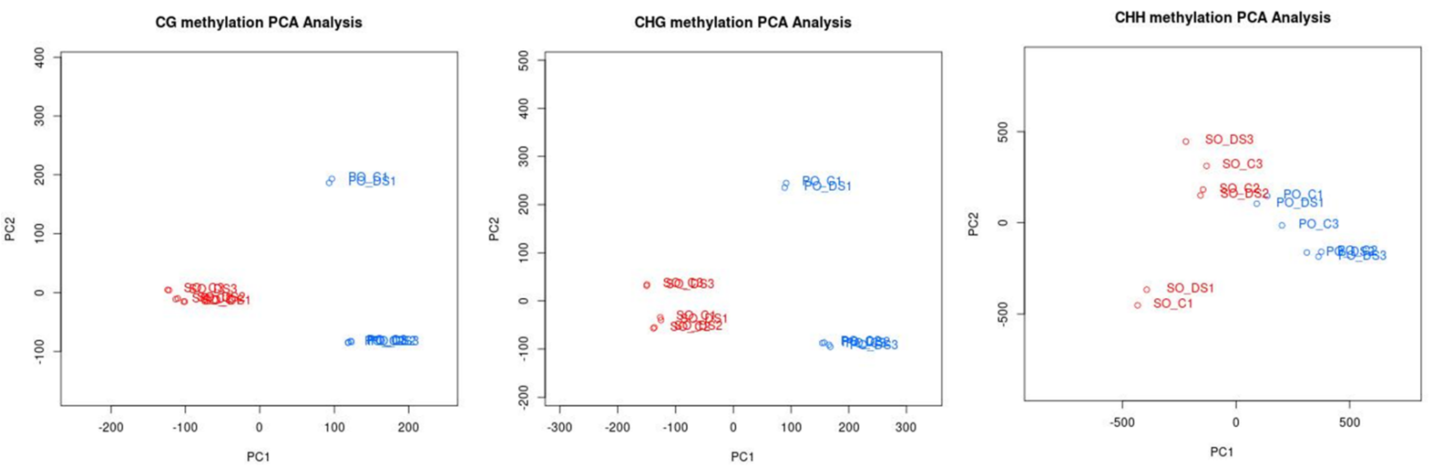


**Fig. S8** Hierarchical clustering analysis based on the percentage methylation profiles of SO (in red) and PO (in blue).
For each species, the three biological replicates for the control and drought stress samples were represented. The method used for hierarchical clustering is 1-Pearson's correlation distance.

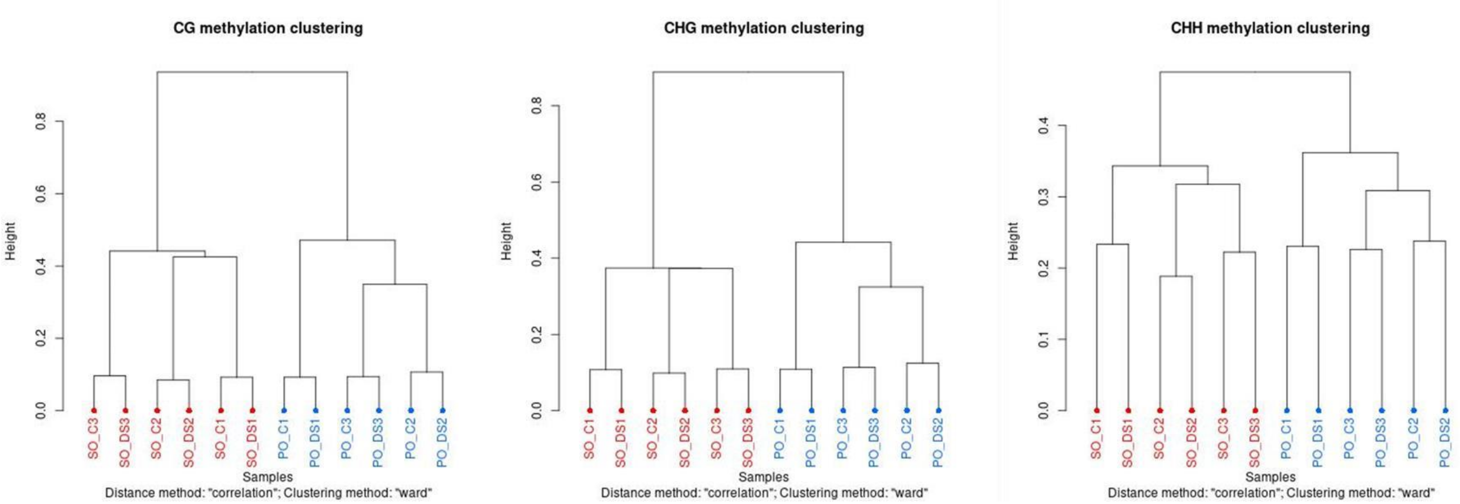

**Fig. S9** Distribution of siRNA (A) and miRNA (B) clusters according to their sequence size from 20-nt to 24-nt.

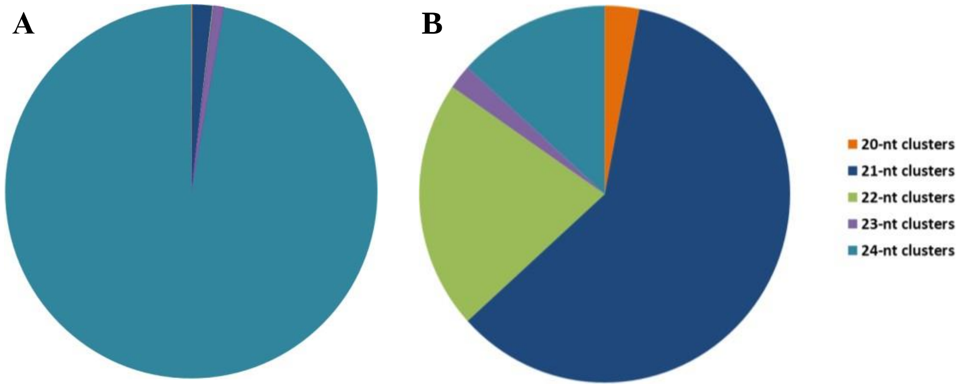


**Fig. S10** Percentages of the small RNA clusters according to their genomic annotation.
Only the smRNA clusters having an annotation on gene bodies and/or 2 kb promoter regions and/or repeated sequences are represented. ‘Gene bodies’, ‘2 kb promoter’ and ‘Repeats’ referred to smRNA clusters annotated only in one of these three categories. Clusters annotated in at least two of these categories were also reported : ‘Gene bodies & Promoter’, ‘Gene bodies & Repeats’ and ‘Promoter & Repeats’. The percentages of each genomic annotation were calculated over the total number of clusters having an annotation.

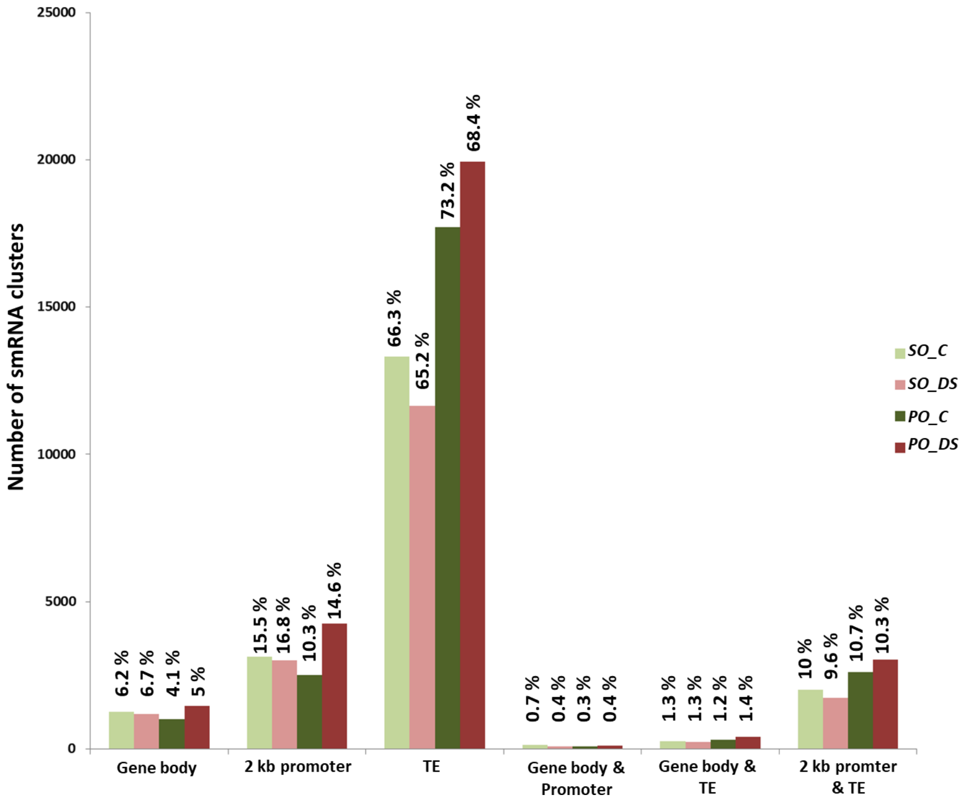

**Fig. S11** Identification of overlaps between 24-nt smRNA clusters and DMRs for PO (C vs. DS), SO (C vs. DS), DS (SO_DS *vs.* PO_DS) and C (SO_C *vs.* PO_C) analyses
(A) General strategy for identification of the most relevant overlaps defined as those with a 24-nt smRNA cluster specific to a single condition and associated to hypermethylation of the genomic region (B) Number of overlaps identified in each analysis for which the presence of a condition-specific 24-nt smRNA cluster is associated with hypermethylation of the region

A)

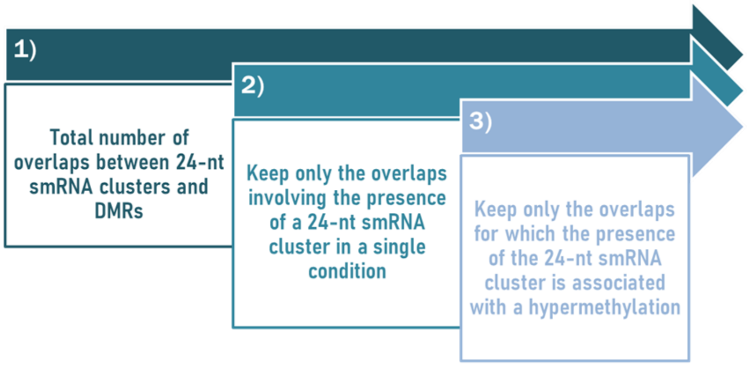


B)

| Analysis | 24-nt smRNA cluster condition | CG | CHG | CHH | Total_Overlaps |
|---|---|---|---|---|---|
| *PO* | PO_C | 17 | 24 | 97 | 138 |
| | PO_DS | 3 | 3 | 97 | 103 |
| *SO* | SO_C | 4 | 7 | 74 | 85 |
| | SO_DS | 3 | 4 | 101 | 108 |
| C | SO_C | 180 | 212 | 315 | 707 |
| | PO_C | 1198 | 1311 | 1162 | 3671 |
| DS | SO_DS | 221 | 219 | 468 | 908 |
| | PO_DS | 818 | 880 | 1025 | 2723 |

**Table S1** Summary of the main metrics of the 12 RNA-seq samples

ME(%) : percentage of mapping efficiency

| Species | Treatment | Samples | Raw Reads | Mapped Reads | ME (%) | % successfully assigned alignments |
|---|---|---|---|---|---|---|
| SO | C | SO_C1 | 36889523 | 27856097 | 75.5 | 75.5 % |
| SO | C | SO_C2 | 28995437 | 22616337 | 78 | 77.4 % |
| SO | C | SO_C3 | 26288753 | 20354757 | 77.4 | 78.5 % |
| PO | C | PO_C1 | 27932560 | 21841211 | 78.2 | 78% |
| PO | C | PO_C2 | 26818372 | 20847454 | 77.7 | 78.3 % |
| PO | C | PO_C3 | 48120961 | 37126044 | 77.2 | 76.9 % |
| SO | DS | SO_DS1 | 35730135 | 28056766 | 78.5 | 78.2 % |
| SO | DS | SO_DS2 | 26064752 | 20396608 | 72.7 | 77.2 % |
| SO | DS | SO_DS3 | 38459786 | 29562514 | 76.9 | 77.5 % |
| PO | DS | PO_DS1 | 49127845 | 38063797 | 77.5 | 77.7 % |
| PO | DS | PO_DS2 | 58218772 | 45761148 | 78.6 | 78.6 % |
| PO | DS | PO_DS3 | 44780143 | 34765438 | 77.6 | 77.6 % |

**Table S3** Gene ontology enrichments for the four gene sets linked to the transcriptome data The GO enrichment analysis were performed on biological process annotations. Only the significant terms were reported according with a p-value <=0.05 (classic column).

| Gene set | GO.ID | Term | Annotated | Significant | DownR | UpR | Expected | p-value |
|---|---|---|---|---|---|---|---|---|
| **gene set #1** | GO:0006952 | defense response | 55 | 4 | 2 | 2 | 0.8 | 0.0083 |
| SO_C vs. SO_DS | GO:0071577 | zinc ion transmembrane transport | 16 | 2 | 0 | 2 | 0.23 | 0.0221 |
| | GO:0048544 | recognition of pollen | 199 | 7 | 3 | 4 | 2.9 | 0.0260 |
| | GO:0006426 | glycyl-tRNA aminoacylation | 2 | 1 | 0 | 1 | 0.03 | 0.0289 |
| | GO:0009733 | response to auxin | 21 | 2 | 1 | 1 | 0.31 | 0.0369 |
| | GO:0006741 | NADP biosynthetic process | 3 | 1 | 0 | 1 | 0.04 | 0.0431 |
| **gene set #2** | GO:0008152 | metabolic process | 6187 | 237 | 152 | 85 | 210.72 | 4.2e-09 |
| PO_C vs. PO_DS | GO:0006468 | protein phosphorylation | 972 | 58 | 47 | 11 | 33.1 | 1.0e-05 |
| | GO:0055114 | oxidation-reduction process | 909 | 51 | 39 | 11 | 30.96 | 0.00017 |
| | GO:0048544 | recognition of pollen | 199 | 13 | 11 | 2 | 6.78 | 0.01825 |
| **gene set #3** | GO:0016998 | cell wall macromolecule catabolic process | 39 | 2 | 2 | 0 | 0.1 | 0.0042 |
| SO_DS vs. PO_DS | GO:0009089 | lysine biosynthetic process via diaminopimelate | 2 | 1 | 1 | 0 | 0.01 | 0.0050 |
| | GO:0006032 | chitin catabolic process | 54 | 2 | 2 | 0 | 0.14 | 0.0079 |
| | GO:0006694 | steroid biosynthetic process | 8 | 1 | 0 | 1 | 0.02 | 0.0199 |
| **gene set #4** | GO:0030244 | cellulose biosynthetic process | 43 | 3 | 0 | 3 | 0.18 | 0.00078 |
| SO_C vs. PO_C | GO:0016998 | cell wall macromolecule catabolic process | 39 | 2 | 2 | 0 | 0.17 | 0.01190 |
| | GO:0006032 | chitin catabolic process | 54 | 2 | 2 | 0 | 0.23 | 0.02208 |
| | GO:0005975 | carbohydrate metabolic process | 401 | 7 | 4 | 3 | 1.71 | 0.02376 |
| | GO:0006098 | pentose-phosphate shunt | 6 | 1 | 0 | 1 | 0.03 | 0.02537 |
| | GO:0032784 | regulation of DNA-templated transcription, elongation | 10 | 1 | 1 | 0 | 0.04 | 0.04194 |

**Table S5** Data description of whole-genome bisulfite sequencing reads for the 3 replicates for each species SO and PO in each condition (control, C and drought stress DS) Total reads : cleanned sequencing reads after data filtrattion ; ME : mapping efficiency ; mC, mCG, mCHG, mCHH percent : percentages of methylated C in each context

| Species | Treatment | Samples | Total reads | % Bisulfite convertion | Mapped reads | ME (%) | Sequencing depth | Total C's analysed | Total C's methylated | mC percent (%) | mCG percent (%) | mCHG percent (%) | mCHH percent (%) |
|---|---|---|---|---|---|---|---|---|---|---|---|---|---|
| **Q. petraea, SO** | **Control, C** | **SO_C1** | 207419840 | 99.4% | 116788505 | 56.3% | 14.8 | 2264088529 | 310127505 | 13.7% | 57% | 37% | 5.6 % |
| **Q. petraea, SO** | **Control, C** | **SO_C2** | 197385968 | 99.4% | 121945514 | 61.8% | 15.5 | 2302200035 | 286451851 | 12.4% | 56.6% | 36.2% | 5.3% |
| **Q. petraea, SO** | **Control, C** | **SO_C3** | 186661756 | 99.4% | 110221842 | 59.1% | 14 | 2095144637 | 293381809 | 14% | 56% | 35.4% | 5% |
| **Q. robur, PO** | **Control, C** | **PO_C1** | 184640778 | 99.4% | 115525998 | 62.6% | 14.7 | 2181647848 | 267750400 | 12.3% | 58% | 36.7% | 5% |
| **Q. robur, PO** | **Control, C** | **PO_C2** | 199333988 | 99% | 112163130 | 56.3% | 14.1 | 2144504771 | 283716455 | 13.2% | 56.6% | 37% | 5.4% |
| **Q. robur, PO** | **Control, C** | **PO_C3** | 203872392 | 99.2% | 128047621 | 62.8% | 16.3 | 2430648244 | 324151770 | 13.3% | 57% | 37.8% | 5% |
| **Q. petraea, SO** | **Drought stress, DS** | **SO_DS1** | 202942956 | 99% | 125295638 | 61.7% | 16 | 2443350528 | 339333880 | 13.9% | 57.5% | 37.1% | 5.8% |
| **Q. petraea, SO** | **Drought stress, DS** | **SO_DS2** | 219392340 | 99.4% | 135962669 | 62% | 18 | 2600797417 | 348607985 | 13.4% | 57.9% | 36.3% | 5.5% |
| **Q. petraea, SO** | **Drought stress, DS** | **SO_DS3** | 182349236 | 99.2% | 111397615 | 61.1% | 14.1 | 2092364764 | 279268652 | 13.3% | 57% | 36% | 5.3% |
| **Q. robur, PO** | **Drought stress, DS** | **PO_DS1** | 220672202 | 99.5% | 136095829 | 61.7% | 17.3 | 2470964957 | 324741660 | 13.1% | 57.3% | 35.7% | 4.6% |
| **Q. robur, PO** | **Drought stress, DS** | **PO_DS2** | 202085600 | 99.2% | 115545879 | 57.2% | 14.5 | 2172574172 | 292828324 | 13.5% | 56.8% | 37.2% | 3.8% |
| **Q. robur, PO** | **Drought stress, DS** | **PO_DS3** | 185291698 | 99% | 115963439 | 62.6% | 14.7 | 2176392762 | 297604390 | 13.7% | 59.1% | 38.1% | 5.4% |

**Table S6** Genomic annotations for DMRs identified between C and DS samples of SO (A) and PO (B)
'Gene body' (intron + exon), '2 kb promoter' and 'TE' referred to DMRs annotated only in one of these three categories. Clusters annotated in at least two of these categories were also reported: 'Gene body & Promoter', 'Gene body & TE' and 'Promoter & TE'. The percentages of each genomic annotation were calculated over the total number of clusters having an annotation

**(A)**

| | CG | CHG | CHH |
|---|---|---|---|
| **Gene body** | 34 (12.9 %) | 53 (6.3 %) | 31 (2.1 %) |
| **2 kb promoter** | 30 (11.4 %) | 50 (5.9 %) | 251 (17.2 %) |
| **TE** | 168 (63.6 %) | 695 (82.3 %) | 933 (63.8 %) |
| **Gene body & Promoter** | 7 (2.7 %) | 6 (0.7 %) | 11 (0.8 %) |
| **Gene body & TE** | 3 (1.1 %) | 3 (0.4 %) | 21 (1.4 %) |
| **Promoter & TE** | 22 (8.3 %) | 37 (4.4 %) | 216 (14.8 %) |
| **Total** | **264** | **844** | **1463** |

**(B)**

| | CG | CHG | CHH |
|---|---|---|---|
| **Gene body** | 20 (7.1 %) | 36 (5.2 %) | 28 (1.5 %) |
| **2 kb promoter** | 27 (9.6 %) | 36 (5.2 %) | 294 (15.6 %) |
| **TE** | 217 (77 %) | 575 (82.7 %) | 1234 (65.4 %) |
| **Gene body & Promoter** | 3 (1.1 %) | 5 (0.7 %) | 12 (0.6 %) |
| **Gene body & TE** | 1 (0.4 %) | 5 (0.7 %) | 33 (1.7 %) |
| **Promoter & TE** | 14 (5 %) | 38 (5.5 %) | 287 (15.2 %) |
| **Total** | **282** | **695** | **1888** |

**Table S7** Genomic annotations for DMRs identified between SO and PO in control (A) and drought stress (B) conditions by distinguishing hypo- and hypermethylated DMRs 'Gene body', '2 kb promoter' and 'TE' referred to DMRs annotated only in one of these three categories. Clusters annotated in at least two of these categories were also reported: 'Gene body & Promoter', 'Gene body & TE' and 'Promoter & TE'. The percentages of each genomic annotation were calculated over the total number of clusters having an annotation.

**(A)**

| | CG | | CHG | | CHH | |
|---|---|---|---|---|---|---|
| | **HypoM** | **HyperM** | **HypoM** | **HyperM** | **HypoM** | **HyperM** |
| **Gene body** | 2052 (22 %) | 1088 (20.5 %) | 914 (8.2 %) | 497 (8.3 %) | 51 (2.6 %) | 73 (2.7 %) |
| **2 kb promoter** | 730 (7.8 %) | 602 (11.3 %) | 664 (5.9 %) | 494 (8.2 %) | 340 (17.3 %) | 358 (13 %) |
| **TE** | 5841 (62.6 %) | 3165 (60 %) | 8724 (78 %) | 4558 (75.8 %) | 1264 (64.4 %) | 1923 (70 %) |
| **Gene body & Promoter** | 191 (2 %) | 128 (2.4 %) | 203 (1.8 %) | 132 (2.2 %) | 9 (0. 5%) | 8 (0.3 %) |
| **Gene body & TE** | 135 (1.4 %) | 80 (1.5 %) | 163 (1.5 %) | 65 (1.1 %) | 22 (1.1 %) | 31 (1.1 %) |
| **Promoter & TE** | 376 (4 %) | 246 (4.6 %) | 525 (4.7 %) | 269 (4.5 %) | 278 (14.2 %) | 355 (13 %) |
| **Total annotated DMRs** | **9325** | **5309** | **11193** | **6015** | **1964** | **2748** |
| **Total DMRs** | **11166** | **16916** | **18534** | **10989** | **3331** | **4429** |
| **Percentage of annotated DMRs** | **83.5 %** | **31.4 %** | **60.4 %** | **54.7 %** | **59%** | **62%** |

**(B)**

| | CG | | CHG | | CHH | |
|---|---|---|---|---|---|---|
| | **HypoM** | **HyperM** | **HypoM** | **HyperM** | **HypoM** | **HyperM** |
| **Gene body** | 1938 (21.9 %) | 1211 (20.2 %) | 905 (9 %) | 568 (8.3 %) | 56 (2.9 %) | 71 (1.9 %) |
| **2 kb promoter** | 685 (7.7 %) | 641 (10.7 %) | 603 (6 %) | 543 (8 %) | 262 (13.7 %) | 436 (11.7 %) |
| **TE** | 5553 (62.6 %) | 3655 (60.8 %) | 7786 (77.1 %) | 5180 (75.8 %) | 1288 (67.3 %) | 2743 (73.6 %) |
| **Gene body & Promoter** | 203 (2.3 %) | 132 (2.2 %) | 207 (2 %) | 139 (2 %) | 11 (0.6 %) | 5 (0.13 %) |
| **Gene body & TE** | 133 (1.5 %) | 97 (1.6 %) | 129 (1.3 %) | 111 (1.6 %) | 25 (1.3 %) | 43 (1.2 %) |
| **Promoter & TE** | 357 (4 %) | 272 (4.5 %) | 471 (4.7 %) | 291 (4.3 %) | 273 (14.3 %) | 430 (11.5 %) |
| **Total annotated DMRs** | **8869** | **6008** | **10101** | **6832** | **1915** | **3728** |
| **Total DMRs** | **16146** | **12039** | **16863** | **12128** | **3254** | **5980** |
| **Percentage of annotated DMRs** | **55%** | **50%** | **60%** | **56%** | **59%** | **62%** |

**Table S8** Characteristics of the 97 DMRs showing differences between SO and PO identified in both conditions C and DS. The methylation of these 97 DMRs changes between the two conditions. The genomic locations of these DMRs were also noted.

| | **Total** | **Gene body** | **2 kb Promoter** | **TE** |
|---|---|---|---|---|
| **DMR hyperM under DS and hypoM in C condition** | 54 | 2 | 7 | 37 |
| **DMR hypoM under DS and hyperM in C condition** | 43 | 1 | 5 | 19 |

**Table S9** Genomic annotations for the 36315 common DMRs identified between SO and PO in C and DS conditions
'Gene body', '2 kb promoter' and 'TE' referred to DMRs annotated only in one of these three categories. Clusters annotated in at least two of these categories were also reported: 'Gene body & Promoter', 'Gene body & TE' and 'Promoter & TE'.

| **Total common DMRs** | 36315 |
|---|---|
| **Common DMRs annotated** | 21037 |
| ***Gene body*** | 1530 |
| ***2 kb promoter*** | 664 |
| ***TE*** | 15760 |
| ***Gene body & Promoter*** | 241 |
| ***Gene body & TE*** | 1760 |
| ***Promoter & TE*** | 1082 |

**Table S10** DMRs between PO and SO identified in both C and DS conditions are enriched in highly differenciated SNPs between SO and PO.
SNPs differentiation between the two species PO and SO were obtained from Leroy et al (2019). Two gene sets corresponding to the common DMRs (i) 'DMRgene' annotated in gene bodies (2699 genes) and (ii) 'DMR2kbpromoter' annotated in the 2 kb promoter region (1531 genes) were tested for enrichments for genes including SNPs with FST values above the 0.9, 0.95, 0.99 and 0.999 quatile of the genome-wide FST distribution, respectively. Significance of enrichement estimtes were assessed using 1000 permutations and exist with a p-value < 0.001

| Gene sets | Ngenes | fst_quantile | prop | mean_rprop | CI_rprop | enrichment | pval |
|---|---|---|---|---|---|---|---|
| DMRgene | 2699 | 0.9 | 0.782512041496851 | 0.67095479807336 | [0.654:0.688] | 1.1662664068337 | <0,001 |
| DMRgene | 2699 | 0.95 | 0.614672100778066 | 0.479319748054835 | [0.461:0.498] | 1.28238426076229 | <0,001 |
| DMRgene | 2699 | 0.99 | 0.205631715450167 | 0.138774731381993 | [0.127:0.152] | 1.48176626538834 | <0,001 |
| DMRgene | 2699 | 0.999 | 0.0340866987773249 | 0.0150496480177844 | [0.011:0.02] | 2.26494990029297 | <0,001 |
| DMR2kbpromoteur | 1531 | 0.9 | 0.763553233180927 | 0.670711952971914 | [0.646:0.695] | 1.13842198547027 | <0,001 |
| DMR2kbpromoteur | 1531 | 0.95 | 0.581972566949706 | 0.478657086871326 | [0.455:0.503] | 1.2158444592426 | <0,001 |
| DMR2kbpromoteur | 1531 | 0.99 | 0.197256694970607 | 0.139096668843893 | [0.123:0.156] | 1.41812666406833 | <0,001 |
| DMR2kbpromoteur | 1531 | 0.999 | 0.030698889614631 | 0.0151045068582626 | [0.01:0.021] | 2.03243243243243 | <0,001 |

**Table S11** Gene ontology enrichments for the DMRs identified between SO and PO commonly or specifically in C and DS conditions. DMRs with a genomic location within (A) gene body and (B) 2 kb promoter region

The GO enrichment analysis were performed on biological process annotations. Only the significative terms were reported according with a p-value <=0.05

**A)**

| Condition | GO.ID | Term | Annotated | Significant | Expected | HypoM | HyperM | p-value |
|---|---|---|---|---|---|---|---|---|
| DMRs commons to C and DS conditions | GO:0006468 | **protein phosphorylation** | 972 | 120 | 78.06 | 73 | 46 | 4,00E-07 |
| | GO:0007165 | **signal transduction** | 462 | 48 | 37.1 | 26 | 21 | 0.0053 |
| | GO:0006422 | aspartyl-tRNA aminoacylation | 2 | 2 | 0.16 | 0 | 1 | 0.0064 |
| | GO:0017038 | protein import | 6 | 3 | 0.48 | 2 | 1 | 0.0086 |
| | GO:0006777 | Mo-molybdopterin cofactor biosynthetic process | 3 | 2 | 0.24 | 2 | 0 | 0.0183 |
| | GO:0009733 | **response to auxin** | 21 | 5 | 1.69 | 5 | 0 | 0.0227 |
| | GO:0005975 | **carbohydrate metabolic process** | 401 | 36 | 32.2 | 26 | 10 | 0.0321 |
| | GO:0006535 | cysteine biosynthetic process from serine | 10 | 3 | 0.8 | 3 | 0 | 0.0404 |
| | GO:0007018 | **microtubule-based movement** | 33 | 6 | 2.65 | 5 | 1 | 0.0449 |
| DMRs specific to C condition | GO:0006281 | **DNA repair** | 78 | 12 | 4.21 | 8 | 4 | 0.0054 |
| | GO:0006886 | **intracellular protein transport** | 138 | 15 | 7.44 | 9 | 6 | 0.0086 |
| | GO:0006468 | **protein phosphorylation** | 972 | 68 | 52.41 | 39 | 29 | 0.0130 |
| | GO:0016192 | **vesicle-mediated transport** | 127 | 13 | 6.85 | 8 | 5 | 0.0217 |
| | GO:0008272 | sulfate transport | 12 | 3 | 0.65 | 1 | 2 | 0.0238 |
| | GO:0006914 | autophagy | 5 | 2 | 0.27 | 2 | 0 | 0.0260 |
| | GO:0006546 | glycine catabolic process | 6 | 2 | 0.32 | 1 | 1 | 0.0376 |
| | GO:0051225 | spindle assembly | 6 | 2 | 0.32 | 1 | 1 | 0.0376 |
| DMRs specific to DS condition | GO:0007165 | **signal transduction** | 462 | 36 | 26.07 | 14 | 22 | 0.0069 |
| | GO:0015969 | guanosine tetraphosphate metabolic process | 3 | 2 | 0.17 | 1 | 1 | 0.0092 |
| | GO:0006430 | lysyl-tRNA aminoacylation | 10 | 3 | 0.56 | 2 | 1 | 0.0159 |
| | GO:0006139 | **nucleobase-containing compound metabolic process** | 1002 | 57 | 56.54 | 32 | 25 | 0.0257 |
| | GO:0006511 | **ubiquitin-dependent protein catabolic process** | 64 | 8 | 3.61 | 4 | 4 | 0.0266 |
| | GO:0019674 | NAD metabolic process | 5 | 2 | 0.28 | 1 | 1 | 0.0284 |
| | GO:0006754 | ATP biosynthetic process | 26 | 3 | 1.47 | 2 | 1 | 0.0284 |
| | GO:0006468 | **protein phosphorylation** | 972 | 68 | 54.85 | 31 | 37 | 0.0329 |
| | GO:0045454 | cell redox homeostasis | 33 | 5 | 1.86 | 3 | 2 | 0.0359 |
| | GO:0000162 | tryptophan biosynthetic process | 6 | 2 | 0.34 | 1 | 1 | 0.0410 |

**B)**

| Condition | GO.ID | Term | Annotated | Significant | Expected | HypoM | HyperM | p-value |
|---|---|---|---|---|---|---|---|---|
| DMRs commons to C and DS conditions | GO:0009733 | **response to auxin** | 21 | 6 | 0.95 | 4 | 2 | 0.00025 |
| | GO:0006468 | **protein phosphorylation** | 972 | 65 | 43.85 | 39 | 26 | 0.00060 |
| | GO:0007165 | **signal transduction** | 462 | 31 | 20.84 | 13 | 18 | 0.01909 |
| | GO:0009231 | riboflavin biosynthetic process | 7 | 2 | 0.32 | 2 | 0 | 0.03667 |
| DMRs specific to C condition | GO:0001522 | **pseudouridine synthesis** | 16 | 5 | 0.7 | 3 | 2 | 0.00044 |
| | GO:0032259 | methylation | 28 | 4 | 1.22 | 3 | 1 | 0.02069 |
| | GO:0006869 | lipid transport | 16 | 3 | 0.7 | 1 | 2 | 0.02995 |
| DMRs specific to DS condition | GO:0071577 | zinc ion transmembrane transport | 16 | 4 | 0.77 | 4 | 0 | 0.0062 |
| | GO:0042026 | protein refolding | 4 | 2 | 0.19 | 1 | 1 | 0.0131 |

**Table S12** Number of overlaps identified between i) DMRs and the gene body of DEGs and ii) DMRs and the 2 kb promoter region of DEGs in the different comparative analyzes

| Analysis for overlap | Nber of DMR in overlap with DEG | | | | Nber of commons overlap |
|---|---|---|---|---|---|
| | CG_context | CHG_context | CHH_context | Total | |
| PO_DMR-DEG_gene body | 4 | 4 | 2 | 10 | 1 |
| *PO*_DMR-DEG_2kb promoter | 4 | 6 | 33 | 43 | |
| *SO*_DMR-DEG_gene body | 2 | 1 | 4 | 7 | 2 |
| *SO*_DMR-DEG_2kb promoter | 0 | 5 | 15 | 20 | |
| DS_DMR-DEG_gene body | 27 | 16 | 2 | 45 | 16 |
| DS_DMR-DEG_2kb promoter | 19 | 14 | 9 | 42 | |
| C_DMR-DEG_gene body | 33 | 23 | 6 | 62 | 17 |
| C_DMR-DEG_2kb promoter | 16 | 21 | 14 | 51 | |

**Table S14** Summary of the main metrics for the twelve small-RNA samples

| Species | Treatment | Samples | Total_reads | Reads_afeter_Rfam_cleaning | % of ME on chloroplastic genome | Reads afetr chloroplast mapping | Reads mappped on ref Q. robur g enome | % of ME |
|---|---|---|---|---|---|---|---|---|
| Q. petraea, SO | Control, C | SO_C1 | 9216673 | 7646495 | 17.09 | 6339674 | 3201816 | 50.5 |
| Q. petraea, SO | Control, C | SO_C2 | 6965134 | 5934734 | 9.81 | 5352541 | 2973426 | 55.55 |
| Q. petraea, SO | Control, C | SO_C3 | 9821103 | 8417719 | 18.92 | 6825400 | 3282851 | 48.1 |
| Q. petraea, SO | Drought stress, DS | SO_DS1 | 8898855 | 7313197 | 19.29 | 5902801 | 2896117 | 49.06 |
| Q. petraea, SO | Drought stress, DS | SO_DS2 | 5480534 | 5000411 | 7.76 | 4612611 | 2552156 | 55.33 |
| Q. petraea, SO | Drought stress, DS | SO_DS3 | 6263909 | 5718369 | 12.94 | 4978560 | 2576995 | 51.76 |
| Q. robur, PO | Control, C | PO_C1 | 5906885 | 5267810 | 14.37 | 4510766 | 2512056 | 55.69 |
| Q. robur, PO | Control, C | PO_C2 | 27081207 | 24424822 | 9.54 | 22095467 | 12115946 | 54.83 |
| Q. robur, PO | Control, C | PO_C3 | 13112025 | 11363401 | 14.23 | 9746946 | 4712822 | 48.35 |
| Q. robur, PO | Drought stress, DS | PO_DS1 | 6410813 | 5856486 | 11.94 | 5157025 | 2854763 | 55.36 |
| Q. robur, PO | Drought stress, DS | PO_DS2 | 14160709 | 13114110 | 5.59 | 12381333 | 6807566 | 54.98 |
| Q. robur, PO | Drought stress, DS | PO_DS3 | 7288577 | 6630469 | 11.7 | 5854899 | 3262282 | 55.72 |

ME : mapping efficiency

**Table S15** Description of the smRNA clusters predicted in SO and PO species in control (C) and drought stress (DS) conditions

| | *SO_C* | *SO_DS* | *PO_C* | *PO_DS* |
|---|---|---|---|---|
| **Total number of clusters** | 37978 | 34660 | 39966 | 52925 |
| **Number of miRNA clusters** | 106 | 88 | 58 | 80 |
| **Number of siRNA clusters** | 37872 | 34572 | 39908 | 52845 |
| **Number of reads per cluster** | 123 | 104 | 100 | 98 |
| **Number of reads per clusters in RPM** | 6.72 | 6.65 | 4.19 | 2.76 |
| **Average cluster lenght (in bp)** | 269 | 272 | 299 | 318 |

rpm : total number of reads per cluster normalized to reads per million
bp : bas paired

**Table S16** Number and percentage of clusters annotated in gene bodies and/or 2 kb promoter and/or repeats for each species (SO and PO) in each condition (C and DS)

| | **SO_C** | **SO_DS** | **PO_C** | **PO_DS** |
|---|---|---|---|---|
| **Total number of clusters** | 37978 | 34660 | 39966 | 52925 |
| **Number of cluster annotated in gene bodies and/or 2 kb promoter region and/or TE** | 20089 | 17860 | 24195 | 29169 |
| **% of clusters with a genomic location** | 52.9 | 51.5 | 60.5 | 55.1 |

**Table S17** Gene ontology enrichments for the common clusters between PO and SO comparative analysis of control (C) vs. drought stress (DS) conditions
The GO enrichment analyses were performed on biological process annotations. Only the significant terms were reported according with a p-value <=0.05

| Condition | GO.ID | Term | Annotated | Significant | Expected | p-value |
|---|---|---|---|---|---|---|
| **PO_C** | GO:0045454 | cell redox homeostasis | 33 | 2 | 0.09 | 0.0037 |
| | GO:0007165 | signal transduction | 462 | 5 | 1.28 | 0.0075 |
| | GO:0006298 | mismatch repair | 13 | 1 | 0.04 | 0.0354 |
| | GO:0006662 | glycerol ether metabolic process | 17 | 1 | 0.05 | 0.0460 |
| **SO_C** | GO:0045454 | cell redox homeostasis | 33 | 2 | 0.1 | 0.0040 |
| | GO:0007165 | signal transduction | 462 | 5 | 1.34 | 0.0091 |
| | GO:0006298 | mismatch repair | 13 | 1 | 0.04 | 0.0370 |
| | GO:0006662 | glycerol ether metabolic process | 17 | 1 | 0.05 | 0.0481 |
| **PO_DS** | GO:0006855 | drug transmembrane transport | 42 | 3 | 0.25 | 0.0019 |
| | GO:0006075 | (1->3)-beta-D-glucan biosynthetic process | 13 | 2 | 0.08 | 0.0026 |
| | GO:0006635 | fatty acid beta-oxidation | 2 | 1 | 0.01 | 0.0118 |
| | GO:0080028 | nitrile biosynthetic process | 5 | 1 | 0.03 | 0.0292 |
| | GO:0019762 | glucosinolate catabolic process | 5 | 1 | 0.03 | 0.0292 |
| | GO:0006952 | defense response | 55 | 2 | 0.32 | 0.0416 |
| | GO:0006289 | nucleotide-excision repair | 8 | 1 | 0.05 | 0.0463 |
| **SO_DS** | GO:0006855 | drug transmembrane transport | 42 | 3 | 0.24 | 0.0017 |
| | GO:0006075 | (1->3)-beta-D-glucan biosynthetic process | 13 | 2 | 0.07 | 0.0023 |
| | GO:0006635 | fatty acid beta-oxidation | 2 | 1 | 0.01 | 0.0113 |
| | GO:0030244 | cellulose biosynthetic process | 43 | 2 | 0.24 | 0.0244 |
| | GO:0080028 | nitrile biosynthetic process | 5 | 1 | 0.03 | 0.0280 |
| | GO:0019762 | glucosinolate catabolic process | 5 | 1 | 0.03 | 0.0280 |
| | GO:0006952 | defense response | 55 | 2 | 0.31 | 0.0384 |
| | GO:0006289 | nucleotide-excision repair | 8 | 1 | 0.05 | 0.0444 |
| | GO:0006099 | tricarboxylic acid cycle | 9 | 1 | 0.05 | 0.0498 |
| | GO:0006505 | GPI anchor metabolic process | 9 | 1 | 0.05 | 0.0498 |

**Table S18** Gene ontology enrichments for the specific clusters of PO in control (C) and drought stress (DS) conditions
The GO enrichment analyses were performed on biological process annotations. Only the significant terms were reported according with a p-value <=0.05

| Condition | GO.ID | Term | Annotated | Significant | Expected | p-value |
|---|---|---|---|---|---|---|
| **PO_C** | GO:0045454 | cell redox homeostasis | 33 | 2 | 0.09 | 0.0037 |
| | GO:0007165 | signal transduction | 462 | 5 | 1.28 | 0.0075 |
| | GO:0006298 | mismatch repair | 13 | 1 | 0.04 | 0.0354 |
| | GO:0006662 | glycerol ether metabolic process | 17 | 1 | 0.05 | 0.0460 |
| **PO_DS** | GO:0055114 | oxidation-reduction process | 909 | 69 | 40.55 | 9.8e-06 |
| | GO:0007165 | signal transduction | 462 | 34 | 20.61 | 6.8e-05 |
| | GO:0006855 | drug transmembrane transport | 42 | 8 | 1.87 | 0.00045 |
| | GO:0006508 | proteolysis | 281 | 20 | 12.54 | 0.00549 |
| | GO:0006094 | gluconeogenesis | 5 | 2 | 0.22 | 0.01815 |
| | GO:0009690 | cytokinin metabolic process | 6 | 2 | 0.27 | 0.02642 |
| | GO:0000162 | tryptophan biosynthetic process | 6 | 2 | 0.27 | 0.02642 |
| | GO:0006310 | DNA recombination | 22 | 4 | 0.98 | 0.03183 |
| | GO:0005975 | carbohydrate metabolic process | 401 | 25 | 17.89 | 0.03453 |

**Table S19** Gene ontology enrichments for the specific clusters of SO in control (C) and drought stress (DS) conditions

The GO enrichment analyses were performed on biological process annotations. Only the significant terms were reported according with a p-value <=0.05

| | GO.ID | Term | Annotated | Significant | Expected | p-value |
|---|---|---|---|---|---|---|
| **SO_C** | GO:0007165 | signal transduction | 462 | 29 | 10.04 | 3.3e-09 |
| | GO:0006855 | drug transmembrane transport | 42 | 5 | 0.91 | 0.002 |
| | GO:0005975 | carbohydrate metabolic process | 401 | 14 | 8.72 | 0.034 |
| | GO:0009813 | flavonoid biosynthetic process | 2 | 1 | 0.04 | 0.043 |
| **SO_DS** | GO:0007165 | signal transduction | 462 | 21 | 6.04 | 5.7e-07 |
| | GO:0006298 | mismatch repair | 13 | 2 | 0.17 | 0.012 |
| | GO:0030244 | cellulose biosynthetic process | 43 | 3 | 0.56 | 0.018 |
| | GO:0006419 | alanyl-tRNA aminoacylation | 3 | 1 | 0.04 | 0.039 |
